# Managing the phyllosphere: Agronomic practices influence the ecology and evolution of *Pseudomonas syringae* in cherry orchards

**DOI:** 10.64898/2026.04.22.719873

**Authors:** Ziyue Zeng, John W. Mansfield, Andrea Vadillo-Dieguez, John Connell, James Irvine, Michelle T. Hulin, Eleftheria Stavridou, Amanda Karlström, Fernando Duarte Frutos, Nastasiya F. Grinberg, Mojgan Rabiey, Richard J. Harrison, Xiangming Xu, Robert W. Jackson

**Affiliations:** Niab, Cambridge, CB3 0LE, UK; School of Biosciences and Birmingham Institute of Forest Research, University of Birmingham, B15 2TT, UK; Faculty of Natural Sciences, Imperial College London, SW7 2AZ, UK; Department of Plant Soil & Microbial Sciences, Michigan State University, MI 48824, USA; Niab at East Malling, Kent, ME19 6BJ, UK

## Abstract

Bacterial canker, caused the *Pseudomonas syringae* species complex, is a major constraint on sweet cherry production worldwide. However, the influence of agronomic practices on pathogen ecology, dispersal and evolution under field conditions remains poorly understood. Here, we combined a factorial-design field experiment with whole-genome sequencing to investigate the effects of polytunnel covering and nitrogen fertigation on phyllosphere populations and the dynamics of a key pathogen, *P. syringae* pathovar *syringae* 9644 (*Pss*9644) in young cherry trees.

Epiphytic *P. syringae* populations initially resembled those in surrounding woodland environments. Over time, pathogenic phylogroup 2d lineages became dominant, particularly on uncovered trees. Diversity of *P. syringae* populations was higher in uncovered treatments. Polytunnel covering markedly altered community composition and limited rain-splash dispersal of *Pss*9644 from stem cankers to leaves, thereby interrupting a key stage of the disease cycle. By contrast, nitrogen fertigation had no detectable effect on phyllosphere community structure, but enhanced plant growth and reduced lesion expansion following inoculation.

Whole-genome sequencing of re-isolated *Pss*9644 strains revealed limited short-term genomic diversification, with single-nucleotide polymorphisms detected in 22 re-isolates. In total, 36 mutations were identified across the chromosome although no mutation affected virulence or motility.

Taken together, our results show that agronomic practices influence both pathogen ecology and disease outcomes through distinct mechanisms: polytunnel covering primarily limits pathogen dispersal and reshapes phyllosphere communities, while nitrogen fertigation enhances plant growth and reduces disease severity. These findings highlight the potential to integrate canopy management and nutrient strategies to mitigate bacterial canker risk in commercial cherry production.

## Introduction

The *Pseudomonas syringae* (*Ps*) species complex is widely distributed in the environment and comprises lineages that include both pathogenic and non-pathogenic strains associated with a diverse range of plant species^1–3^. As one of the most significant bacterial pathogens of fruit trees, *Ps* causes bacterial canker of cherry – a major disease responsible for significant yield losses of this important horticultural crop worldwide^4,5^. Members of numerous *Ps* phylogroups (PGs) are common components of the leaf surface microbiome, where non-pathogenic and potentially pathogenic populations coexist on leaf surfaces^6,7^. The importance of an epiphytic phase in the development of bacterial diseases caused by *Ps* was first highlighted by Crosse^8^ in his pioneering research on bacterial canker of cherry. His analysis showed that the pathogen *Ps* pv. *morsprunorum* Race 1 (in PG3, now designated as *P. amygdali*^9^) was a frequent coloniser of cherry leaf surfaces. He concluded that most leaf-scar infections of cherry trees were caused by epiphytic bacteria washed from the leaf surfaces via rain splash dispersal, but there was no correlation between numbers of epiphytic bacteria present and the frequency of leaf spot symptoms^10^. The epiphytic multiplication of the canker pathogen without causing symptoms was, therefore, identified as a feature of the disease cycle leading to the economically damaging production of cankers due to invasion of woody tissues in the autumn^4,11,12^. A schematic of the cycle is shown in **Figure S1**. In recent surveys of leaf surface colonising bacteria throughout UK orchards and woodlands, epiphytic *Ps* strains with pathogenic potential have been identified including members of PGs 2 and 10^6,7^. These findings highlight the potential for epiphytic populations of *Ps* to impact crop health and losses due to bacterial disease.

Leaf surfaces represent open microbial habitats exposed to continual immigration and emigration of bacterial populations via aerosols and windblown droplets generated by rain splash falling onto mucoid colonies^1^. Crosse^10^ demonstrated that prior to the senescence of overwintered canker lesions, *Ps* is dispersed onto newly emerging leaves and sometimes cause a necrotic leaf spotting, with severity largely depending on the amount of rain during this period. Modern cherry production systems frequently employ polyethylene covers (polytunnels) with the aims of protecting fruit from cracking caused by rainfall and improving fruit quality^13–15^. Beyond the intended purposes, these structures substantially alter the canopy microclimate by reducing rainfall exposure and leaf wetness while modifying humidity, light, and temperature^16,17^, all of which could influence the composition, persistence and spread of epiphytic *Ps* populations.

Irrigation with nitrogen fertiliser (fertigation) is another agronomic practice that can increase cherry fruit yield and has also been proposed as partially responsible for reducing susceptibility to bacterial canker in certain stone fruits, including peach, French prune^18,19^ and almond^20^. However, the influence of covers or fertigation on development of bacterial canker of cherry has not been examined^21^.

Strains of *Ps* are frequently found outside agricultural environments but there has been little analysis of the impact of agronomic practices on the diversity of *Ps* on crop plants. One study of the leaf surface microbiome using 16S RNA gene sequencing has partly addressed this issue in field-grown canola (oilseed rape), *Phaseolus* bean and soybean^22^, however none of these crops was covered or treated with fertilisers. Fertigation would be expected to alter the availability of nutrients within treated plants and might lead to the release of different compounds into the phyllosphere, thereby influencing the composition of leaf surface microflora. Indeed, nutrient management has been shown to reshape phyllosphere community structure and pathogen interactions in other systems. For example, the addition of fertiliser (N:P:K 24-8-16) abolished the microbiome-mediated protection of tomato leaves against infection by *Ps* pv. *tomato*^23^. These findings suggest that agricultural inputs can modify both nutrient fluxes and microbial ecology at the leaf surface, potentially altering the balance between commensal and pathogenic *Ps* populations.

Whole genome sequencing (WGS), as used to examine the diversity of *Ps* in the phyllosphere by Zeng et al.^6^, also allows genomic surveillance of the presence of certain strains of bacteria using stringent cut-offs of average nucleotide identity (ANI). Beyond plant pathology, genomic surveillance has been successfully used to trace the spread of disease from infections of animal pathogens^24–26^ and provides more selectivity than using PCR-based diagnostics^25^. The growing use of WGS to examine the evolution and epidemiology of bacterial pathogens of plants has been recently reviewed by Straub et al.^27^, highlighting its value for understanding pathogen transmission and adaptation. In this study, we used WGS to follow the dispersal of a widespread virulent PG2d strain, *Ps* pv. *syringae* 9644 (*Pss*9644), from inoculation sites in the stems of cherry saplings onto the leaf surface, where it persisted epiphytically without visible symptoms. This approach overcame the need to use a strain tagged through genetic modification, for example through the insertion of genes for antibiotic resistance, which is strictly regulated in the UK under the Genetically Modified Organisms (Contained Use) Regulations 2014.

Our multifactorial experiment during the 2022 and 2023 seasons combined visual assessment of canker lesion development with WGS of *Ps* strains recovered from leaf washings. The factorial design allowed us to address the following topics.

1. The establishment of epiphytic *Ps* on newly planted saplings.
2. The impact of covers and nitrogen fertigation on the presence and diversity of *Ps* populations in the phyllosphere.
3. The spread of *Ps* from established cankers to the leaf surface.
4. Genome modifications occurring in the inoculated strain following its spread to the leaf surface.
5. The Influence of agronomic practices and pathogen inoculation on plant growth.

Our results provide answers to each of these topics and support the design of such an experiment combining multiple controlled variables. One of the most significant findings from our study is that polythene covers minimise rain-splash mediated dispersal of bacterial inoculum onto the leaf from woody cankers and thereby lead to a reduction in spread of *Ps* from infected tissues. We confirm, therefore, that the use of covers has the unplanned, beneficial effect of suppressing disease development.

## Results

### The diversity of *P. syringae* phylogroups recovered from the leaves of young cherry trees

The schematic (**Figure 1**) illustrates the full factorial split plot experimental design, with three factors: polythene cover (yes or no), nitrogen fertigation (high or low) and inoculation with a known sweet cherry pathogen *Pss*9644 (yes or no). For each treatment (namely, combination of three factors at a specific level), the same trees were repeatedly sampled for symptomless, visually healthy leaves, from which bacterial colonies with classic *Ps* colony morphology were recovered through leaf washing. In total, 1920 individual bacterial strains were isolated and screened to identify *Ps* via PCR. A subset of 369 PCR-confirmed *Ps* strains were whole genome sequenced. Genomic sequences of 242 of the 369 strains shared at least 95% ANI with a known *Ps* sequence in GenBank^28^, confirming their identity as members of the *Ps* species complex.

**Figure 1.**
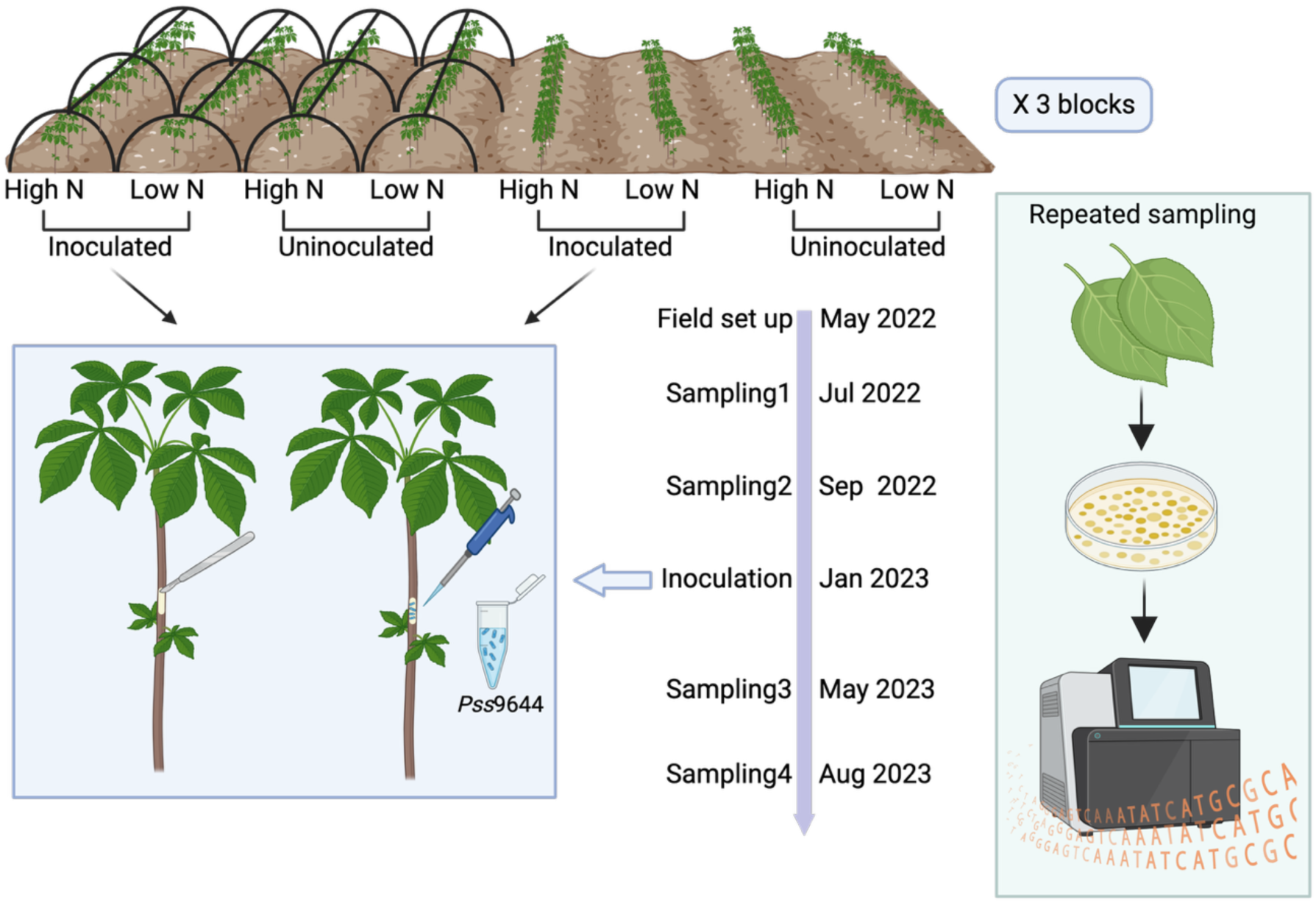
Schematic showing layout of a single block and the sampling timeline. In spring 2022, 240 cherry saplings were planted at NIAB EMR (Kent, UK) in a split-plot design combining polythene cover, nitrogen fertigation, and inoculation with *Pss*9644. Symptomless leaf samples were collected in summer and autumn 2022 and 2023, with inoculation carried out in January 2023. There were ten trees per treatment in each of the three block replicates. The layout of one block is shown here. Leaves of five trees per treatment per block were sampled each time. *Ps* colonies recovered from leaf washes were screened and sequenced.

Data from the analysis of the diversity of *Ps* recovered throughout the sampling experiment are presented as components of the phylogenetic tree in **Figure 2**. Members of PGs 1, 2, 3, 4, 7, 9, 10 and 13 were identified with 90% ANI. Species were further discriminated at 95% ANI. Overall, the more common species recovered were PG2b&d and PG7b (**Table 1**). To achieve finer resolution within dominant lineages, consistent with our previous study^6^, the ANI cut-off was raised to 96%. Under this threshold, the species PG1a&b and PG2b&d were further separated into distinct clades; PG1a and 1b, and 2b and 2d, respectively. *Ps* classification at the level of clade was used in the following analysis. Importantly, the use of WGS allowed the differentiation of the inoculated *Pss*9644 strain from other PG2d lineages that were found at the trial site. The re-isolated *Pss*9644 strains shared over 99.95% ANI with the inoculated strain and are omitted from **Figure 2**, **Table 1**, **Figure 4** and **Figure S4** as the legends specify.

**Figure 2.**
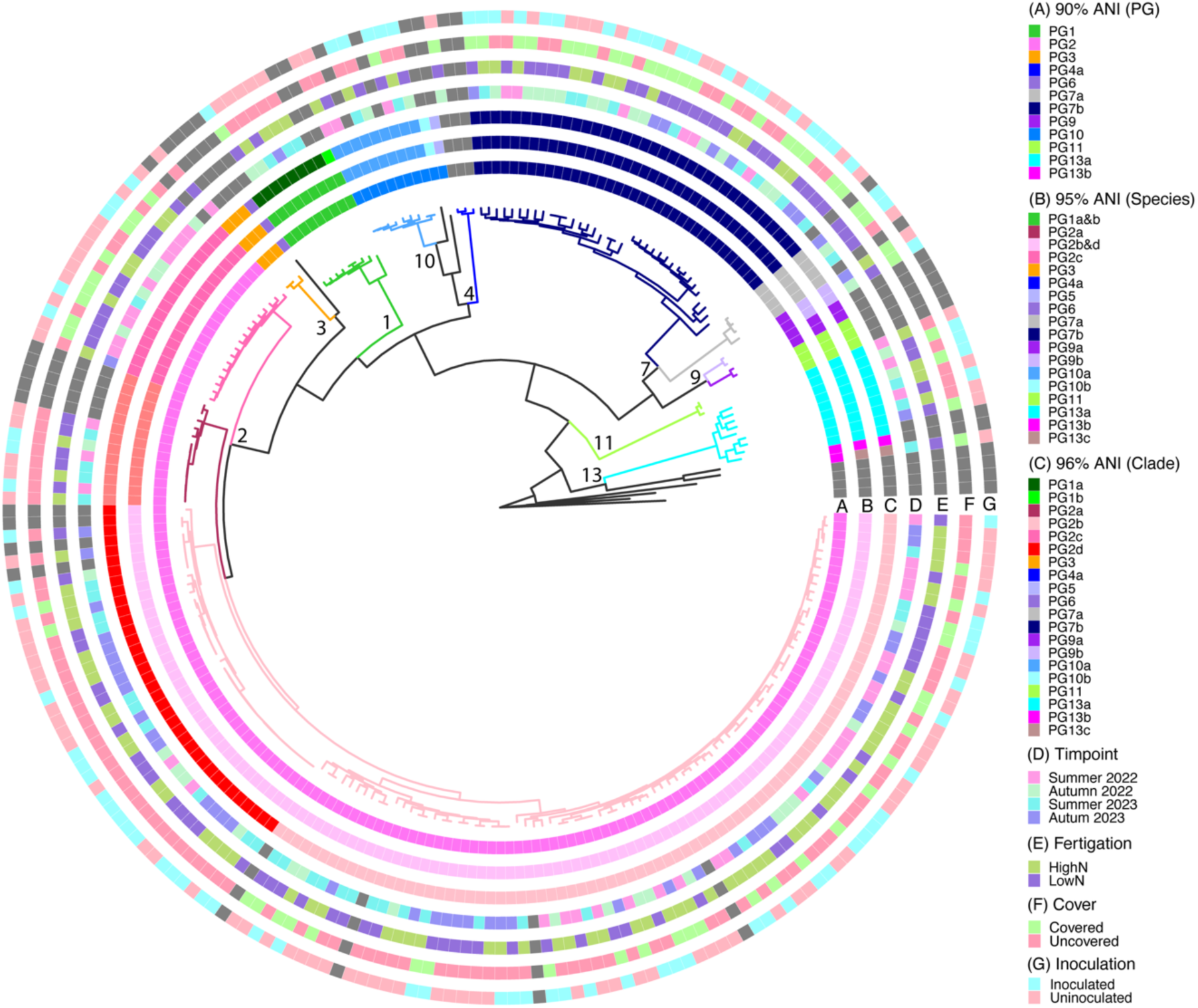
Phylogroups (PGs) of epiphytic *P. syringae* strains recovered from the field experiment at NIAB East Malling in 2022 and 2023. A maximum likelihood core genome phylogeny tree showing *Ps* strains isolated from the field and 39 reference strains. Rings A, B and C show classifications of *Ps* strains at an ANI of 90%, 95% and 96% respectively. Rings D – G show sampling time points and specific treatments. Reference strains are marked in dark grey in rings D-G. Numbers near branches show PG classification. Colours of tree branches show classification of species. Forty-two *Ps*s9644-like strains, the progeny of inoculum, were excluded from this figure.

**Figure 3.**
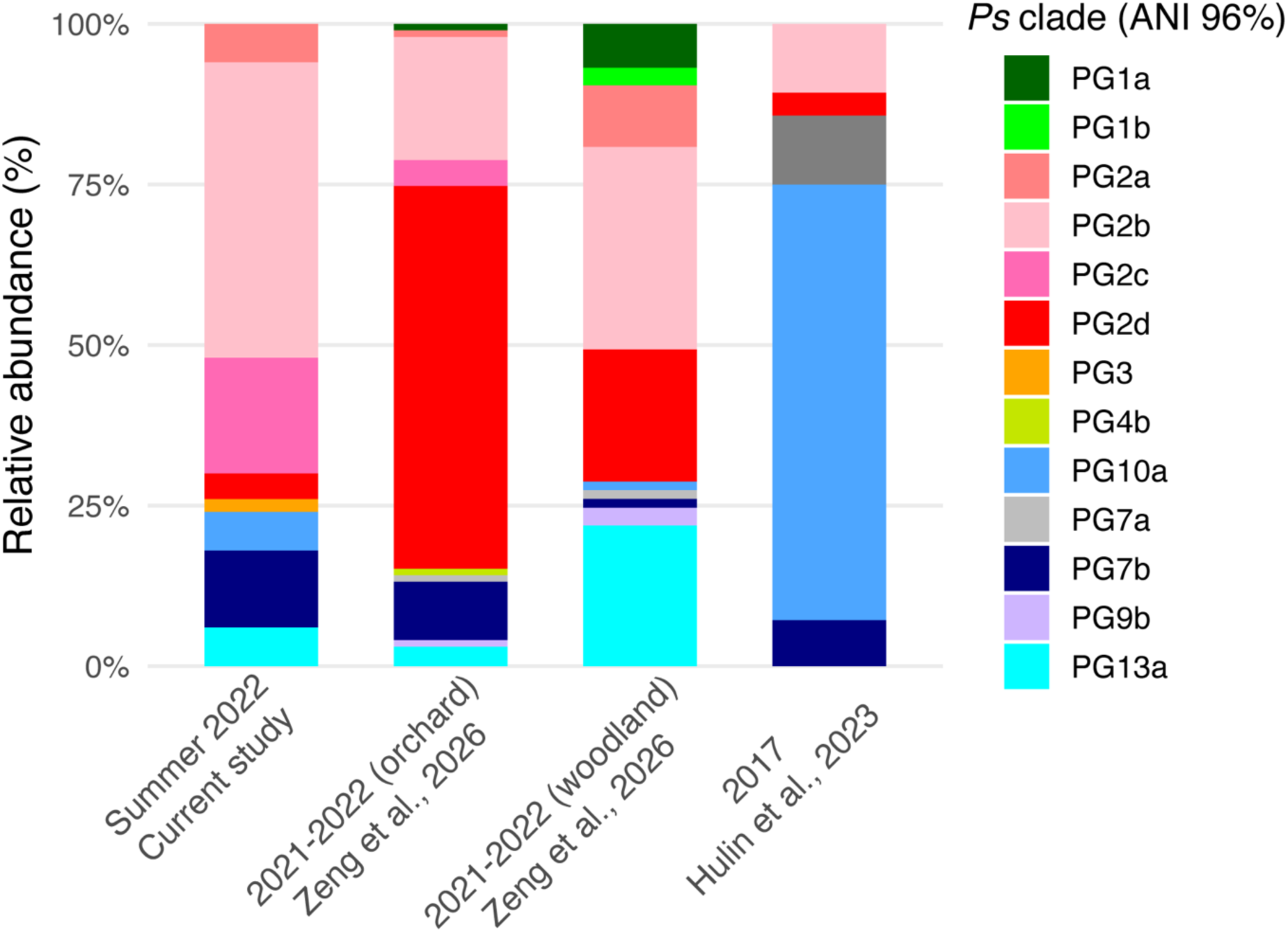
Comparison of *P. syringae* epiphytic populations recovered from different sampling experiments. Relative abundances of clades (based on 96% ANI clustering) are shown for isolates collected from the first sampling of the current study in summer 2022, commercial cherry orchards and nearby woodlands in southeast UK sampled in 2021–2022 (combined data presented), and an earlier survey in 2017. Bars represent the proportional composition of phylogroups within each dataset.

**Figure 4.**
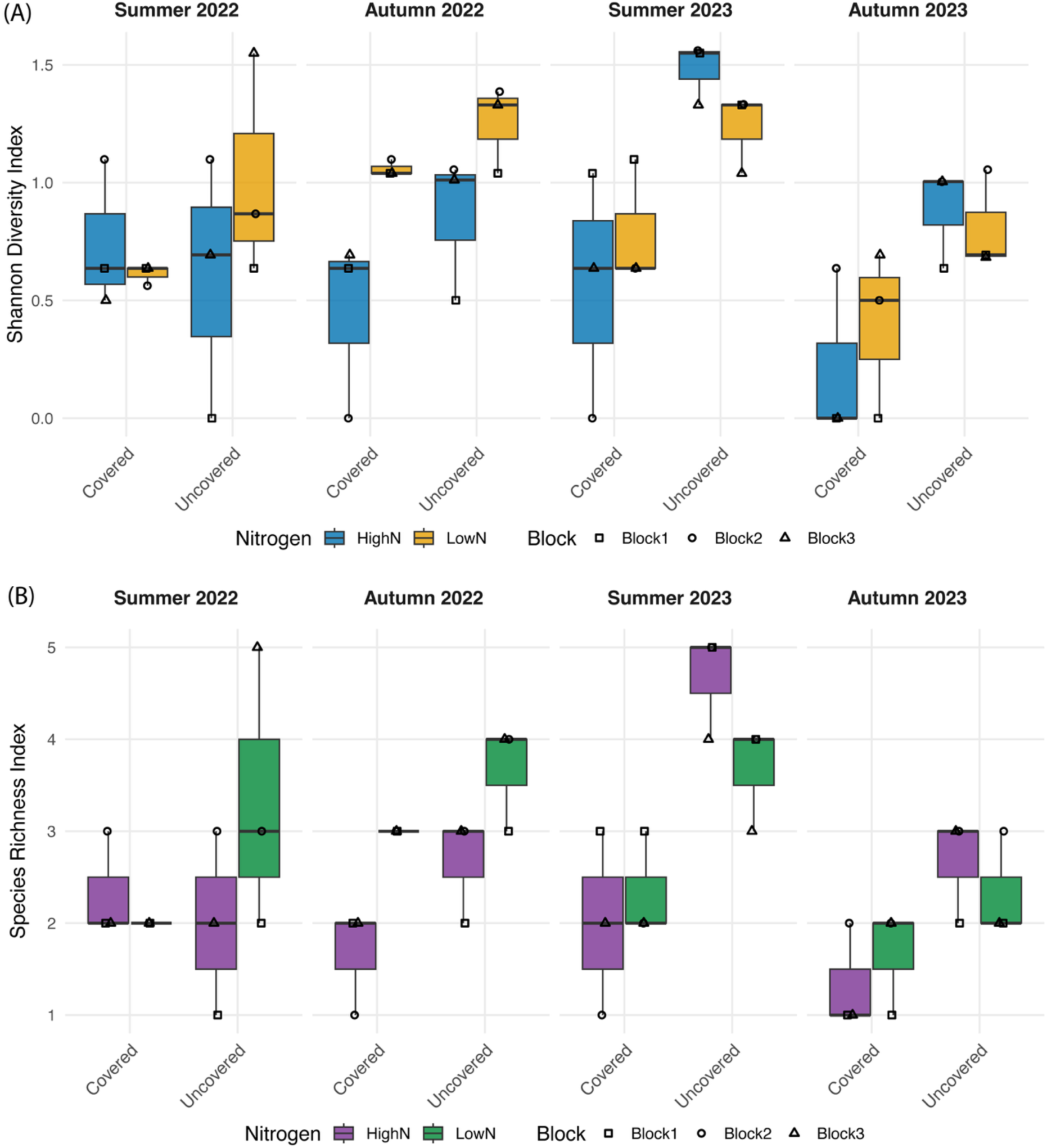
Effects of cover and nitrogen treatments on P*. syringae* diversity and species richness across timepoints. Boxplots show **(A)** Shannon diversity and **(B)** species richness of *Ps* populations associated with cherry leaves in the covered and uncovered groups, across high– and low-levels of nitrogen fertigation. Each point represents an individual block replicate, with the median shown as a horizontal line. Forty-two *Ps*s9644-like strains (ANI ≥ 99.95%), progeny of inoculum, were excluded.

**Table 1.** Numbers of, and percentages of different phylogroups (PGs), species and clades amongst the *P. syringae* strains recovered from four samplings. The *Ps* strains recovered were classified into PGs, species and clades based on 90%, 95% and 96% average nucleotide identities (ANIs). Results are marked in blue if they were the same as those at PG level (ANI=90%). Forty-two *Ps*s9644-like strains (ANI ≥ 99.95%), progeny of inoculum, were excluded.

| PG and No. of strains | Percent | Species and No. of strains | Percent | Clades and No. of strains | Percent |
| --- | --- | --- | --- | --- | --- |
| PG1(6) | 3.0% | PG1a&b(6) | 3.0% | PG1a(6) | 3.0% |
| PG2(136) | 68.0% | PG2a(8) | 4.0% | PG2a(8) | 4.0% |
|  |  | PG2b&d(112) | 56.0 % | PG2b(79) | 39.5% |
|  |  |  |  | PG2d(33) | 16.5% |
|  |  | PG2c(16) | 8.0% | PG2c(16) | 8.0% |
| PG3(1) | 0.5% | PG3(1) | 0.5% | PG3(1) | 0.5% |
| PG4a*(1) | 0.5% | PG4a*(1) | 0.5% | PG4a*(1) | 0.5% |
| PG7a*(3) | 1.5% | PG7a*(3) | 1.5% | PG7a*(3) | 1.5% |
| PG7b*(35) | 17.5% | PG7b*(35) | 17.5% | PG7b*(35) | 17.5% |
| PG9(2) | 1.0% | PG9b* (2) | 1.0% | PG9b*(2) | 1.0% |
| PG10(7) | 3.5% | PG10a(7) | 3.5% | PG10a(7) | 3.5% |
| PG13a*(8) | 4.0% | PG13a* (8) | 4.0% | PG13a* (8) | 4.0% |
| PG13b*(1) | 0.5% | PG13b*(1) | 0.5% | PG13b*(1) | 0.5% |

\* Groupings defined from this study based on ANIs and different from reference strains, which are representatives of established clades.

The first sampling of the epiphytic populations of *Ps* in the young trees was carried out at the beginning of the application of various treatments and therefore allowed a comparison with earlier work that focused on established, mature trees at least four years old in commercial cherry orchards, and also on trees within nearby established woodlands^6,7^. The diversity of *Ps* clades was similar to those reported on mature trees in 2021 and 2022, with strains of PG2 predominating. However, most strains recovered from the young trees were PG2b and PG2c as occurred within woodlands whilst PG2d was dominant in the mature trees in orchards in 2021 and 2022. The frequency of occurrence of different PGs altered significantly from 2017 to 2021 and 2022. Compared with the sampling in 2017, PG10a were distinctly under-represented in the more recent surveys (**Figure 3**). During the trial period, a clear increase in frequency of PG2d lineages was recorded in plots without polythene covers even without the inoculated PG2d strain *Pss*9644 (**Figure S2**).

### Impacts of nitrogen, covers and fertigation on the diversity of *P. syringae*

Leaf nutrient analyses confirmed that nitrogen levels were significantly higher in trees that received high-nitrogen fertigation than those in the low-nitrogen groups (p < 0.001; **Figure S3**). Leaf nitrogen concentration also varied significantly across time points (p < 0.001), reflecting the progressive impact of fertigation treatments as the experiment progressed, as well as tree development. Although the focus of this study was on nitrogen, several other nutrients showed less consistent variation associated with time or agronomic treatments (**Figure S3**).

Incorporating nitrogen treatments within the cover analysis revealed that uncovered plants consistently supported higher within-sample (alpha) diversity (**Figure 4**). Cover had a strong effect on both Shannon diversity (p < 0.001) and species richness (p < 0.001), whereas nitrogen showed no significant effects (p = 0.07 for Shannon index, p = 0.11 for richness). Sampling times had moderate effects for both metrics (p = 0.013 for Shannon index, p = 0.003 for richness), reflecting temporal variation in *Ps* populations, but no significant interactions between time and agronomy practices were detected. Principal component analysis supported the impact of covers (**Figure S4**). Samples from uncovered plants clustered separately from those obtained under covers. By contrast, nitrogen fertigation level did not lead to distinct clustering. Consistent with these patterns, PERMANOVA confirmed significant effects of both cover (R² = 0.11, p = 0.001) and timepoint (R² = 0.15, p = 0.009) on community composition, while nitrogen fertigation had no significant effect (R² = 0.01, p = 0.64).

The recovery of *Ps* from covered and uncovered plants in 2022 and 2023 given in **Figure S2** further clarifies the population structure underlying the differences recorded in the alpha diversity with and without covers. Strains of PG2a were only found on uncovered plants. PG2d lineages not associated with the inoculated *Ps* strain were far more frequent on uncovered plants. By contrast, PG13c strains were only found on covered plants and PGs 7a and 7b were more frequent under covers.

### Spread of the inoculated strain *P. syringae* PG2d *Pss*9644

To investigate the impacts of cover and nitrogen fertigation on *Ps* dispersal from lesions on woody tissue to the leaf surface, half of trees were inoculated with *Pss*9644 in January 2023 (**Figure 1**). The inoculated isolate was differentiated from other strains within PG2d by sharing at least 99.95% of ANI with the inoculum *Ps*s9644. No spread of *Ps*s9644 was recorded to any uninoculated plants despite their close proximity to trees with lesions in the plot (**Table S1**). However, spread onto the leaf surface was frequent on trees inoculated in the main stem, particularly on uncovered plants (**Table S1**). No isolates of *Ps*s9644 were recovered from trees under covers in the autumn of 2023 despite artificial inoculation causing canker development (**Table S1**). Spread of inoculated *Pss*9644 onto leaves occurred with very few examples of macroscopically visible symptoms. Some shothole symptoms were seen close to canker lesions (**Figure S5**) but leaves showing these typical symptoms were not included in the survey of epiphytes.

### Progeny of *Pss*9644 on the leaf surface exhibit genetic variation without loss of virulence

Genome sequencing of 42 *Pss*9644-like strains isolated from the leaf surface revealed variations among their genomes. Examination of fine-scale sequence variations identified single-nucleotide polymorphisms (SNPs) in 22 re-isolated strains of *Pss*9644 using Snippy, with the complete long-read genome of *Pss*9644 (GenBank accession GCA_023277945.1) as the reference. In total, 36 variants (SNPs and small deletions) were detected across the 6.2 Mb chromosome, representing 28 unique genomic locations (**Figure S6**). No variations were observed in plasmids.

Several SNPs occurred within coding regions, leading to changes in predicted amino acids in genes encoding products such as, pectate lyase, chemotaxis protein (*cheA*) and DNA repair ATPase (**Table 2**). A distinct hotspot of variation was observed at position 5,383,744, corresponding to a small deletion within the intergenic region between *erpA* and N-acetyl-gamma-glutamyl-phosphate reductase. The deletion was found in seven strains, and six of them carried an identical 7-bp deletion corresponding to a direct-repeat tract (GCTACAA), while one strain exhibited a larger deletion spanning seven repeat units (5,383,744 – 5,383,793) within a region of 16 contiguous direct repeats. Apart from the localized deletion hotspot, no clustering of SNPs was observed across the genome.

**Table 2.**
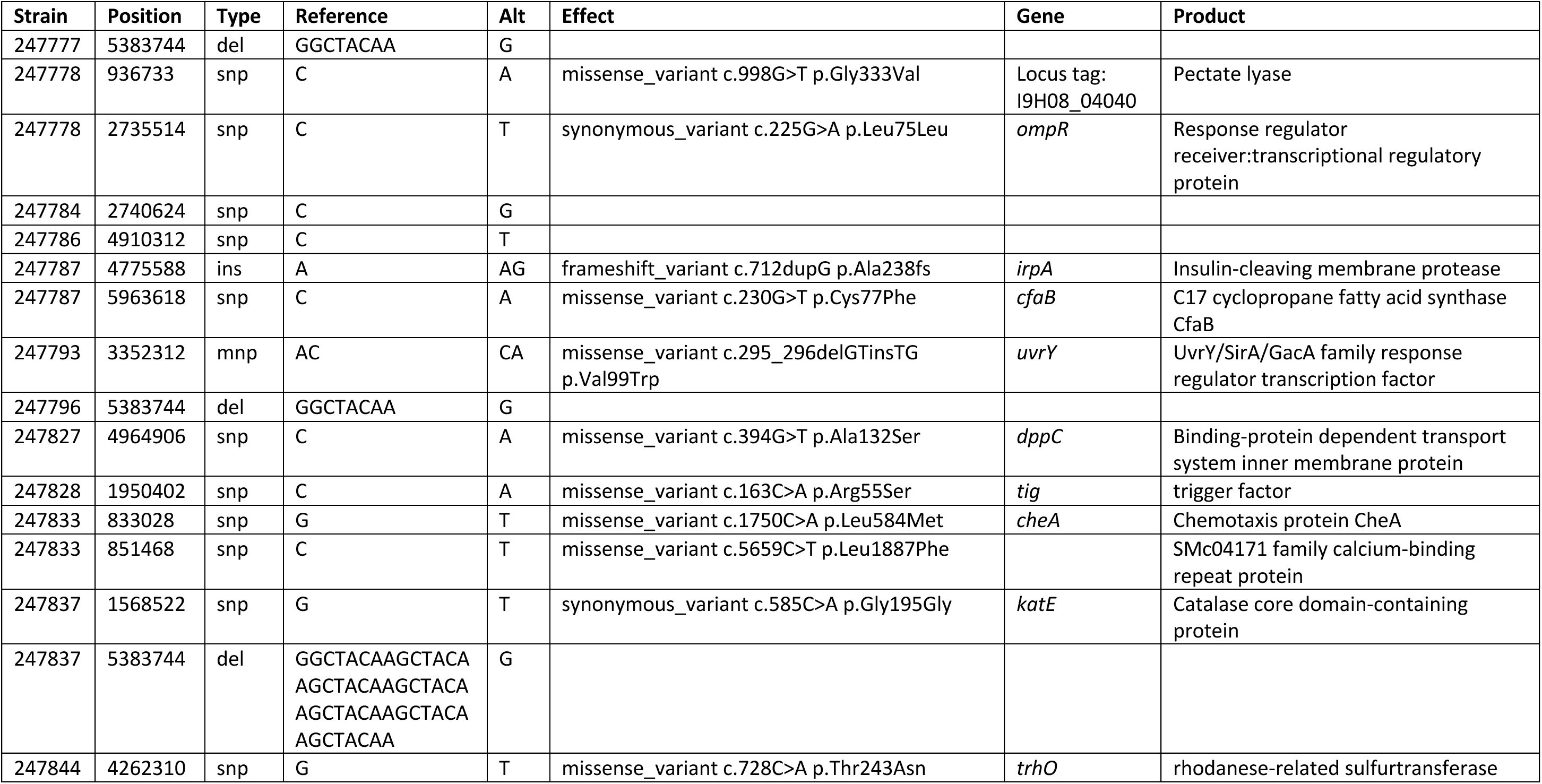
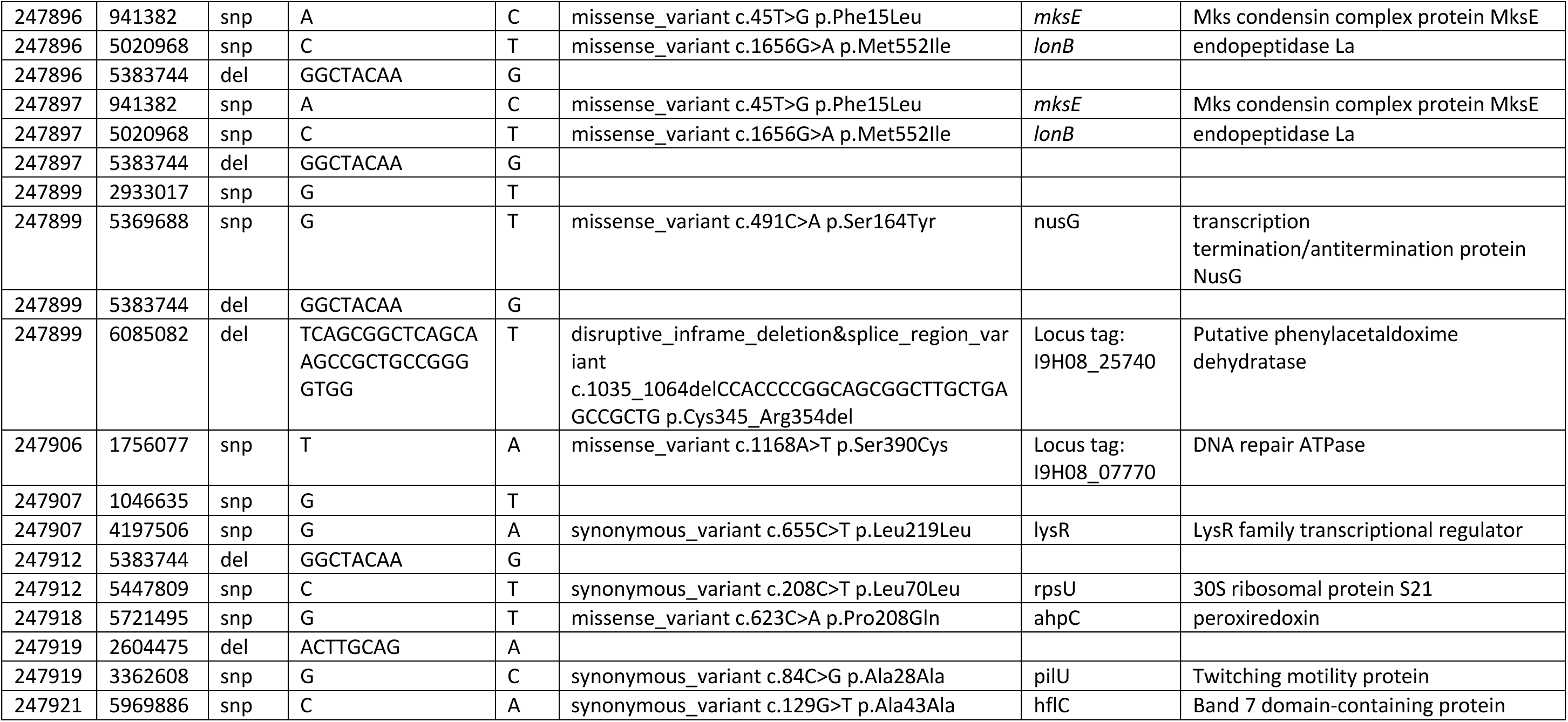
Summary of single-nucleotide polymorphisms (SNPs) and small indels identified in re-isolated *P. syringae* pv. *syringae* 9644 (*Pss*9644) strains relative to the reference genome. Variants were called using Snippy. Columns show the strain identifier, chromosomal position, variant type (snp, single-nucleotide polymorphism; del, deletion; ins, insertion; mnp, multi-nucleotide polymorphism), reference and alternate (Alt) alleles, predicted mutation effect, affected gene and corresponding function annotation.

To test whether genomic variations identified among re-isolated strains of *Pss*9644 affected virulence in cherry, a pathogenicity test was performed using a leaf infiltration assay. The re-isolated strains produced symptoms comparable to those caused by *Pss*9644, with similar degrees of tissue necrosis and discolouration observed (**Figure S7**). Statistical analyses (Kruskal–Wallis test, p = 0.064; Dunn’s post hoc tests, all adjusted p = 1) confirmed that no significant differences in virulence were detected among the strains six days post-inoculation. These results indicated that the genomic variations accumulated during the experiment did not measurably alter pathogenicity under the tested conditions. Similarly, no change in swimming or swarming was observed in the variant with missense mutation in the chemotaxis protein (**Figure S8**).

### Impact of management practices on plant growth and susceptibility to *Ps*s9644

To investigate the effect of cover and nitrogen treatments on lesion development, lesion length was measured six months after pathogen inoculation (**Figure 5**). Lesions were significantly longer on trees fertigated with the low level of nitrogen compared with the high level of nitrogen (p = 0.009), whereas covers had no significant effect on lesion expansion (p = 0.49).

**Figure 5.**
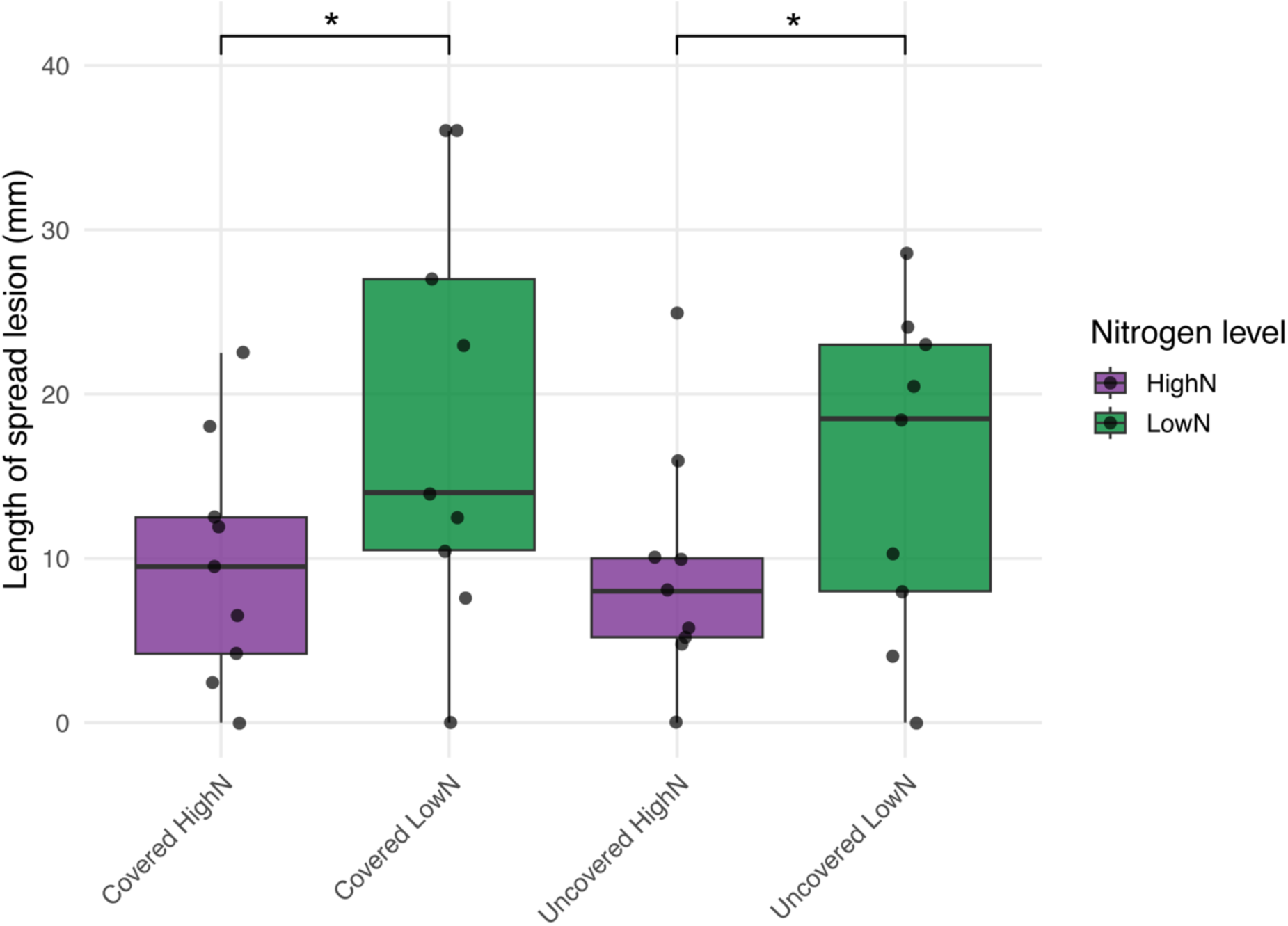
Effect of cover and nitrogen treatments on lesion development following inoculation. Boxplots show the spread lesion length recorded six months post-inoculation. To quantify lesion spread, the surface bark was removed and lesion extension both upwards and downwards from the inoculation site was measured and summed. Boxes indicate the interquartile range, horizontal lines represent medians, and whiskers extend to 1.5× the interquartile range. Points show individual tree measurements. The asterisk (*) indicates a statistically significant difference (p < 0.05).

Although the frequency of *Ps* recovery from leaf surfaces was not significantly influenced by fertigation treatment, increased nitrogen input did affect plant growth and lesion formation after inoculation. As shown in **Figures 6 (A)**, trunk diameters measured at the beginning of the experiment (June 2022) were similar across all treatments. By July 2023, trees fertigated with the high level of nitrogen showed greater trunk diameters compared with the low level of nitrogen (p < 0.001). Neither cover nor inoculation influenced trunk diameter, and no interactions between treatment factors were detected. Baseline height in June 2022 differed among nitrogen treatments (p = 0.015), with trees in the high nitrogen group being shorter at the start of the experiment (**Figure 6 (B)**). Analysis of height increment from June 2022 to July 2023 showed that these initially shorter trees showed greater height gain. Moreover, nitrogen and cover both significantly promoted height growth (p = 0.002 and p = 0.017, respectively), whereas inoculation had no significant effect (p = 0.27). No interactions were statistically significant.

**Figure 6.**
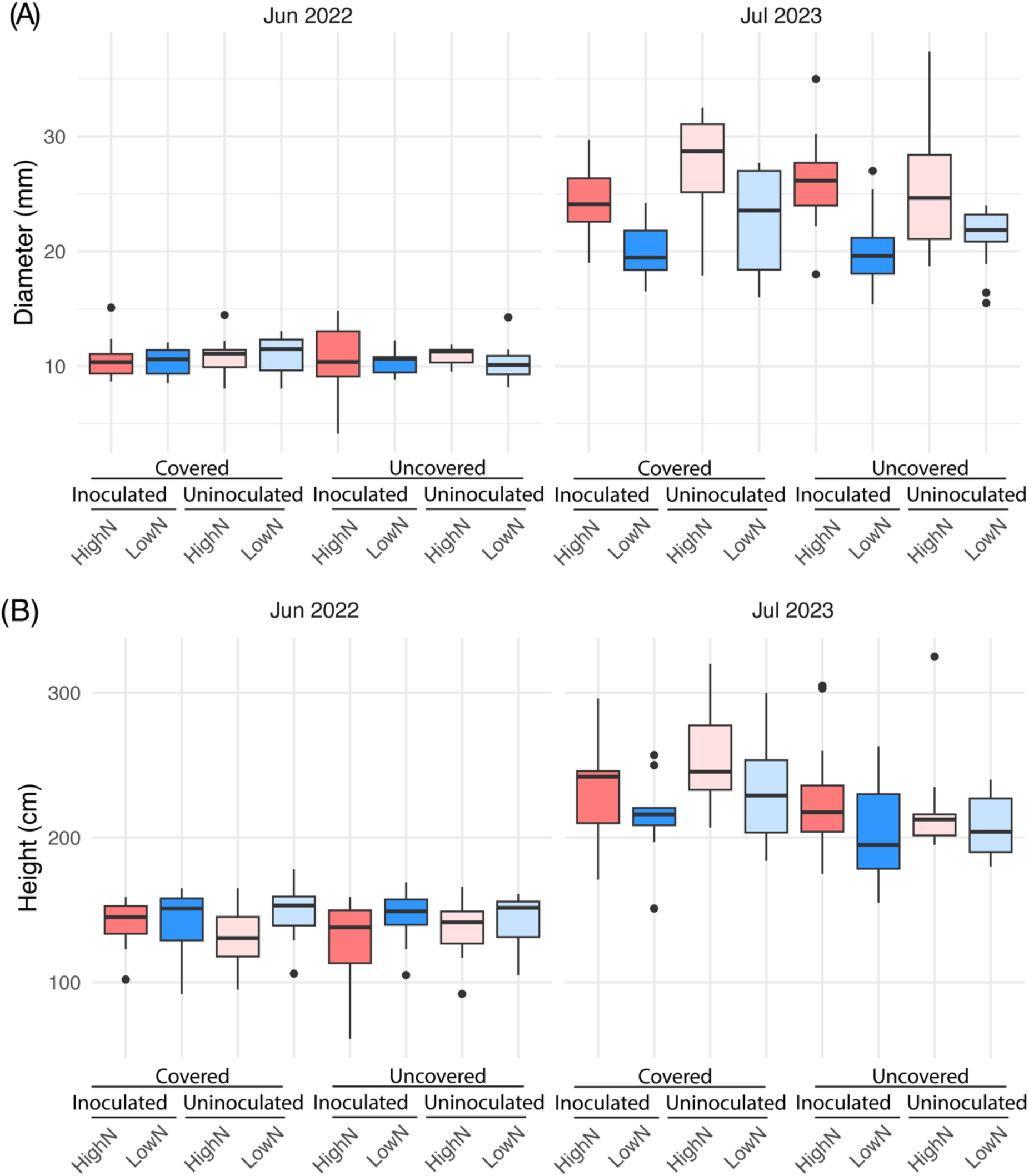
Measurement of (A) trunk diameter and (B) height in June 2022 and July 2023 under different cover, nitrogen, and inoculation treatments. The measurement was performed using a measuring tape at the first and the last timepoints. For each treatment, four trees in each block were measured, three blocks in total. Each box plot shows the measurements of 12 trees. The measurements in July, October in 2022 and January, April, May, June in 2023 are not shown here.

## Discussion

Our results indicate that the epiphytic populations of *Ps* on the young trees used in the experimental plots were initially closely comparable to those from woodland^6^, implying likelihood of bacterial invasion from the wider environment. In particular, a much lower frequency of PG2d strains, the major pathogenic lineage within orchards in the Southeast of England, was observed in the first season. The subsequent, rapid accumulation of PG2d in samples of the leaf surface has not been associated with an increased frequency of shot hole symptoms in leaves or cankers within woody tissues^6^. The PG2d strains appear highly adapted to colonisation of the leaf surface establishing rapidly on uncovered plants and represent a major threat to future cherry tree health in the UK.

The use of covers had clear effects on epiphyte diversity and species richness, particularly the presence at high frequency of certain strains of PG on uncovered rather than covered plants. The more rapid invasion of uncovered plants by PG2d illustrated the protective effect of covers against environmental inoculum in addition to limited dispersal by rain splash of inoculated *Pss*9644 from stem cankers. By contrast, strains of PG7b were far more frequent under covers. Phylogroup 7 strains often include species like *Pseudomonas viridiflava* and are frequently identified in environmental samples rather than causing major agricultural epidemics^29^. However, lineages of *P. viridiflava* have been reported as pathogens of several plants and also as a rare cause of apricot and cherry canker in France^30,31^. The prevalence of PG7 strains as epiphytes may have been promoted by the higher and more stable temperatures occurring under covers and also as a response to higher relative humidities compared with uncovered plants^16^. The predominance of PG2d did not appear to influence the frequency of detection of members of other PGs, suggesting that there is little competition for favourable niches within the phyllosphere.

In contrast to covers, nitrogen fertigation had no significant effect on *Ps* composition in phyllosphere colonisation. However, it did reduce the spread of cankers after artificial inoculation of stems and promoted plant growth both in height and diameter. Nitrogen availability has been reported to have diverse effects on plant physiology and plant associated microbiomes^18,20^, highlighting the complexity of nitrogen–pathogen interactions. In this study, increased nitrogen supply may lead to the enhancement of plant metabolism, which can prime cherry to respond to invasion by the more rapid activation of plant defences^32^. The mechanisms of resistance operating in cherry have not been investigated, but the generation of antimicrobial conditions through enhanced secondary metabolite biosynthesis may lead to phytoalexin accumulation (*Xanthomonas malvacearum*^33^) and may also promote cell wall modifications that are known to restrict invading bacteria in other plants^34,35^.

The identification of the inoculated strain *Pss*9644 by WGS in those strains recovered from cherry phyllosphere suggested a limited spread of the pathogen to leaf surfaces from canker lesions in the stems of saplings, despite significant rainfall during the experimental period.

Nevertheless, *Pss*9644 did establish well on leaves close to inoculation sites. The “life-cycle” of the cherry canker pathogens has largely been deduced by detailed observations and controlled inoculation experiments completed by Crosse and colleagues^11,12^. Using WGS to analyse spread over several seasons should allow experimental confirmation of proposed infection routes, and movement of inoculum from leaves to woody tissue leading to the emergence of cankers in the following spring (**Figure S1**).

The use of WGS enabled the identification of fine-scale genomic variation within the inoculated strain *Pss*9644 following colonisation of the leaf surfaces. Mutation rates in *Ps* have been studied in *Ps.* pv. *actinidiae* based on the recovery of bacteria from lesions^36,37^. Our analysis of epiphytic strains indicates that numerous small changes in the genome can occur without compromising the ability of variants to colonise the leaf surface or act as pathogens invading cherry leaves. Plasmids in *Ps* are highly dynamic and often associated with horizontal gene transfer and adaptive traits^38^, but no mutation was detected in the plasmids of *Pss*9644 during the present study period. Analyses over additional seasons will be required to investigate if the observed mutations are random or play a role in overwintering or long-term adaptation. Longer-term experiments may allow the selection of variants with enhanced virulence or epiphytic fitness. Similar patterns of limited genomic diversification have been observed in animal-associated bacterial pathogens, such as *Burkholderia dolosa*^39^ and *Staphylococcus aureus*^40^, where small numbers of SNPs accumulate during infection without clear evidence of positive selection. In addition, the repeated deletions observed at a single intergenic locus are consistent with instability of short direct repeats, likely arising through replication slippage, a common source of microvariation in bacterial genomes^41,42^

## Conclusion

In conclusion, this study examined whether and if so, how agronomic practices influence both host resistance/tolerance to *Ps* canker development and the ecology of *Ps* in young cherry trees under field conditions. Using a factorial design combining polytunnel covering, nitrogen fertigation, and inoculation with a known cherry pathogen, we found that both polythene covering and increased nitrogen input promoted tree growth, while increased nitrogen fertigation also reduced the severity of canker lesions. WGS of isolates recovered from leaf surfaces revealed diverse epiphytic *Ps* populations dominated by PG2 lineages, with greater phylogenetic diversity observed on uncovered trees. Sequencing enabled tracking of the pathogen’s movement from woody tissues to the phyllosphere and revealed limited but detectable genomic variation within the inoculated strain during the study period. Overall, these results highlight the close links between agronomic management, canopy microbiology and pathogen evolution in perennial fruit crops.

## Materials and Methods

### Plant materials

Cherry cultivar Sweetheart was grafted with rootstock Gisela 5 at NIAB East Malling, UK in March 2021 and stored in the cold room until being potted with slow-release fertiliser in April 2021. The trees were stored under the polytunnel until planting in on May 5^th^, 2022.

### Field design and layout

To investigate the impacts of the application of polytunnel cover, varied levels of nitrogen in fertigation and the inoculation of cherry pathogen on *Ps* populations, pathogen dispersal and canker disease development, a field of cherry trees was set up at NIAB East Malling, UK. As shown in **Table S2**, a split-plot full factorial design was used in this study. There were two levels of each factor (covered vs uncovered polytunnel, low vs high level of nitrogen fertigation, cherry pathogen inoculated vs uninoculated groups), leading to eight types of combined treatments. The treatments were repeated and randomised in three blocks. Due to the structure of the polytunnels, four rows of covered trees were next to each other within each block and were separated from the uncovered rows by guard trees to minimise edge effects. Two rows of trees inoculated with pathogens were next to each other within each subplot and were separated from the uninoculated rows by guard trees to avoid cross-contamination. In total, there were 43 rows, with 4 m between each row. Ten trees were planted in each row, with 2 m between two trees. The first two and the last two trees of each row were used as guard trees to avoid edge effect.

### Agronomic practice and pathogen inoculation

Trees were covered and fertigated in line with standard agronomic practice between April and October. Composition of nutrients used for fertigation are shown in **Table S3**. The recipe was updated in 2023 to increase the difference between two levels of nitrogen. Soil and leaf samples of different treatment groups were collected for nutrient analyses before and after the application of treatments.

Cherry pathogen *Ps* pathovar *syringae* (*Ps*s9644,) was inoculated in January 2023, following the methods used in Hulin et al.^43^. *Ps*s9644 was inoculated in Lysogeny Broth (LB) and incubated on a shaker overnight at 28 ℃. For the trees in the inoculation groups (A, B, E, F), after centrifugation, bacterial pellets were resuspended in 10 mM MgCl_2_ to a final concentration of 2 × 10^7^ CFU/ml. Bark surface of the tree to be inoculated was sterilised with 70% ethanol, and a shallow wound of 2 cm was made into the trunk using a sterile scalpel. Fifty μl of bacterial culture was pipetted into the wound. The inoculation site was then covered with Parafilm and duct tape.

### Tree condition scoring

The height and trunk diameter (10cm above the grafting point) of four trees per treatment per block were measured monthly after planting. In July 2023, surface bark surrounding inoculation sites was removed using a sterile scalpel, and the length of (spreading) lesion was measured using a tape measure.

### Sample collection

Leaf samples were collected at each of the four timepoints. For each treatment in each block, five trees were randomly selected out of the six experimental trees in each row, with the first two and the last two trees of each row used as guard trees. The same trees were sampled repeatedly at each timepoint. From each tree, two symptomless leaves were collected for bacterial isolation. Each leaf sample was rolled up carefully and kept in a 15 ml Falcon tube. All the samples were stored in a pre-chilled cool box or in a cold room at 4 ℃ before being processed. Bacterial isolation was performed within 24 h post sample collection.

### Bacterial isolation

Epiphytes were isolated, purified, screened following the methods of Zeng et al.^6^.

### Whole genome sequencing

For each combined treatment in each of the three blocks, five PCR-confirmed candidate *Ps* strains were selected for WGS (**Table S4)**. Bacterial genomes were sequenced following the methods of Zeng et al.^6^. The strains sequenced in this study are listed in **Table S5 (Excel file)**, and their sequence reads have been deposited in NCBI Sequence Read Archive under BioProject PRJNA1347059.

### Bioinformatics

Genome assembly and annotation, identification of *Ps* strains, phylogenetics and pangenome analyses, identification of virulence-related genes were performed following the methods of Zeng et al.^6^.

### Microbial diversity analysis

Community data aggregated at the *Ps* clade level were used to generate a sample-by-taxon matrix. Alpha diversity metrics, including Shannon diversity and observed richness, were calculated using the R package vegan. Effects of cover, nitrogen treatment and sampling timepoint on alpha diversity were tested using linear mixed-effects models implemented in the R package lmerTest, with cover, nitrogen, timepoint, and their interactions as fixed effects and block as a random effect.

For community composition analysis, the abundance matrix was Hellinger-transformed and analysed by principal component analysis (PCA) using rda in the R package vegan.

Differences in community composition among treatments were further tested using PERMANOVA (adonis2) with cover, nitrogen, and timepoint included as explanatory variables. Alpha diversity and PCA results were visualised using boxplots and ordination plots in R.

### Pathogenicity tests

The reisolated Pss9644 strains were selected for pathogenicity tests. A detached leaf infiltration assay was performed on cultivated cherry (cv. Sweetheart) following the methods of Hulin et al.^43^. Freshly picked leaves were infiltrated with bacterial suspension (2×10^8^ CFU/ml). The cherry pathogen *Pss*9644 and 10mM MgCl_2_ were used as positive and negative controls, respectively.

### Swimming and swarming motility assays

Swimming and swarming motility of bacterial strains were assessed using soft agar assays. For swimming motility, bacteria were grown overnight on King’s B^44^ (KB) agar plates at 28 °C. Single colonies were inoculated by vertical stabbing into the centre of soft agar plates (0.25% agar in 10% KB broth) and incubated at 28 ℃ for 24 h. For swarming motility, overnight cultures grown in liquid KB broth were adjusted to OD_600_= 1.0 in PBS. A 3 µl aliquot was spotted onto the centre of semi-solid agar plates (0.5% agar in KB broth) and incubated at 28 ℃ for 24h.

Images were captured at 15 h and 23 h post-inoculation for swimming assays, and at 16 h and 24 h post-inoculation for swarming assays. Motility was quantified by measuring the diameter of the bacterial expansion zone from digital images using ImageJ. All assays were performed with at least three independent biological replicates. Differences in motility between strains were analysed using one-way analysis of variance (ANOVA) on mean motility values per strain. Pairwise comparisons were performed using Tukey’s honestly significant difference (HSD) test in R.

## Acknowledgements

The authors thank assistance for short-term technical support from Joshua Weblin, Maite Elisa Rekalde, Siyu Miao, Yuan Xue, Lateefat Anike Arogundade, Maialen Garmendia Calvo, Robert Watts, Ainhoa Ozamiz Barrio, Freya Dover, Adam Donaldson, Sharon Halmkan and Marzena Lipska. We acknowledge the support from NIAB EMR Farm Managers Luis Felgueiras and Graham Caspell. The authors gratefully acknowledge Tally Wright and Ian Mackay for their advice on statistical aspects of the field experimental design, and Katherine G. Hinton for sharing scripts and for helpful discussions on pangenome analysis. Our work was supported by the Bacterial Plant Disease Programme BB/T010746/1 and the JABBS Foundation. The authors also thank the British Society for Plant Pathology for Undergraduate Vacation Bursaries, the Novia Salcedo Fundación foundation, European Master Program in Plant Breeding, the Zero Gravity Foundation and the Hardy Plant Society for funding internships. The authors acknowledge Research Computing at the James Hutton Institute for providing computational resources and technical support for the UKCropDiversity-HPC (BBSRC grants BB/S019669/1 and BB/X019683/1), which contributed to the results presented in this paper.

## Author contributions

ZZ, JWM, MTH, RJH, XX and RWJ designed the study. ZZ, AVD, JI, FDF, JWM, ES and AK performed field samplings and lab experiments. ZZ and JC performed bioinformatics analyses. NFG performed statistical analyses. ZZ and JWM drafted the manuscript. All authors contributed to the interpretation of results and preparation of the article and have approved the final manuscript.

## Data Availability Statement

The datasets supporting the conclusions of this article are included within the article and the Supporting Information. The sequences obtained in this study have been submitted to NCBI Sequence Read Archive under BioProject PRJNA1347059.

## Supplementary information

Supplementary material is available as separate files online.

---

**Figure S1.**
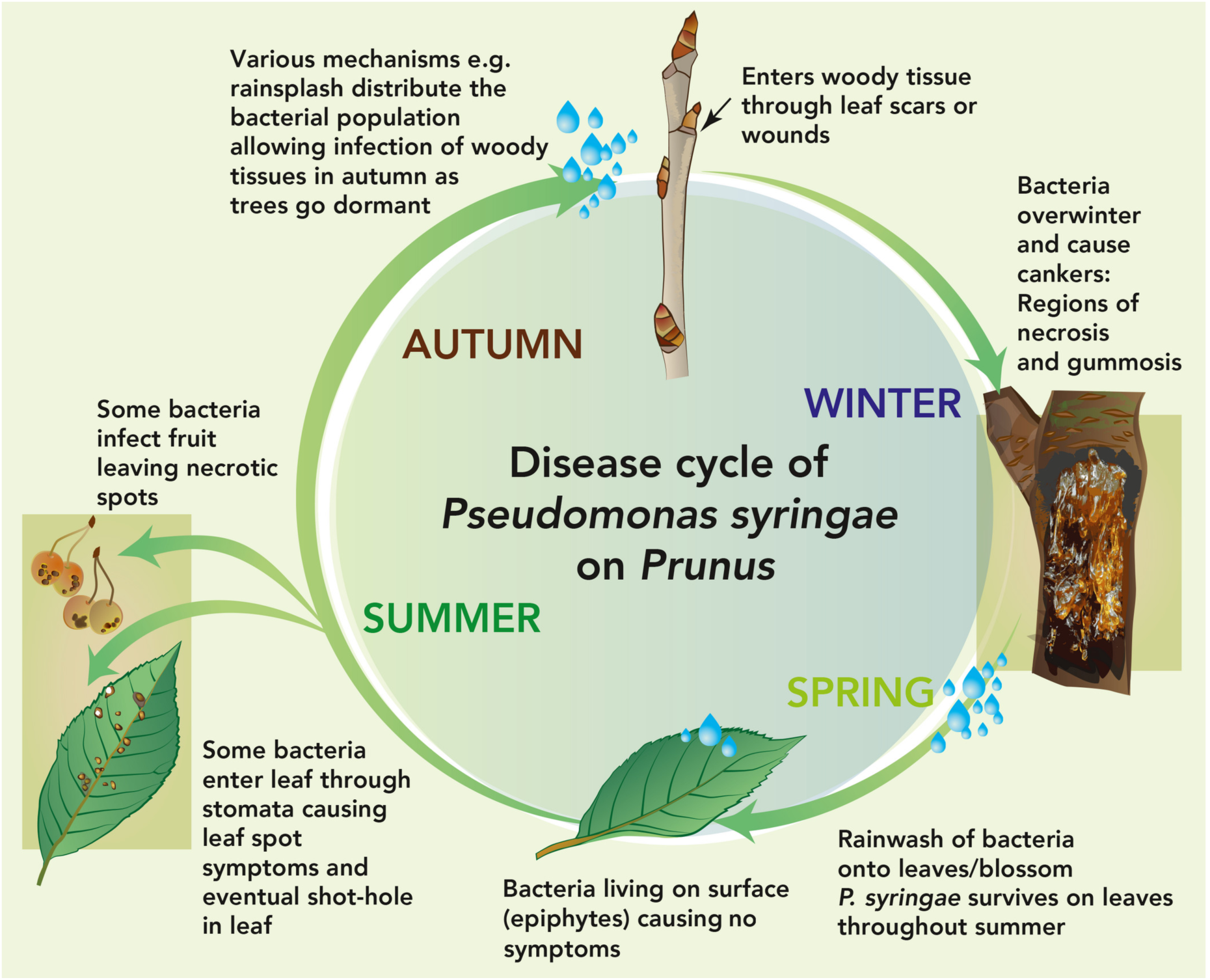
Overview of the canker disease cycle on *Prunus*. Pathogens occupy multiple niches on and within the host at different stages of the disease cycle. Figure reproduced from Hulin et al.^1^, Figure 1. 1 Hulin MT, Jackson RW, Harrison RJ, Mansfield JW. Cherry picking by pseudomonads: After a century of research on canker, genomics provides insights into the evolution of pathogenicity towards stone fruits. *Plant Pathol* 2020; **69**: 962–978.

**Figure S2.**
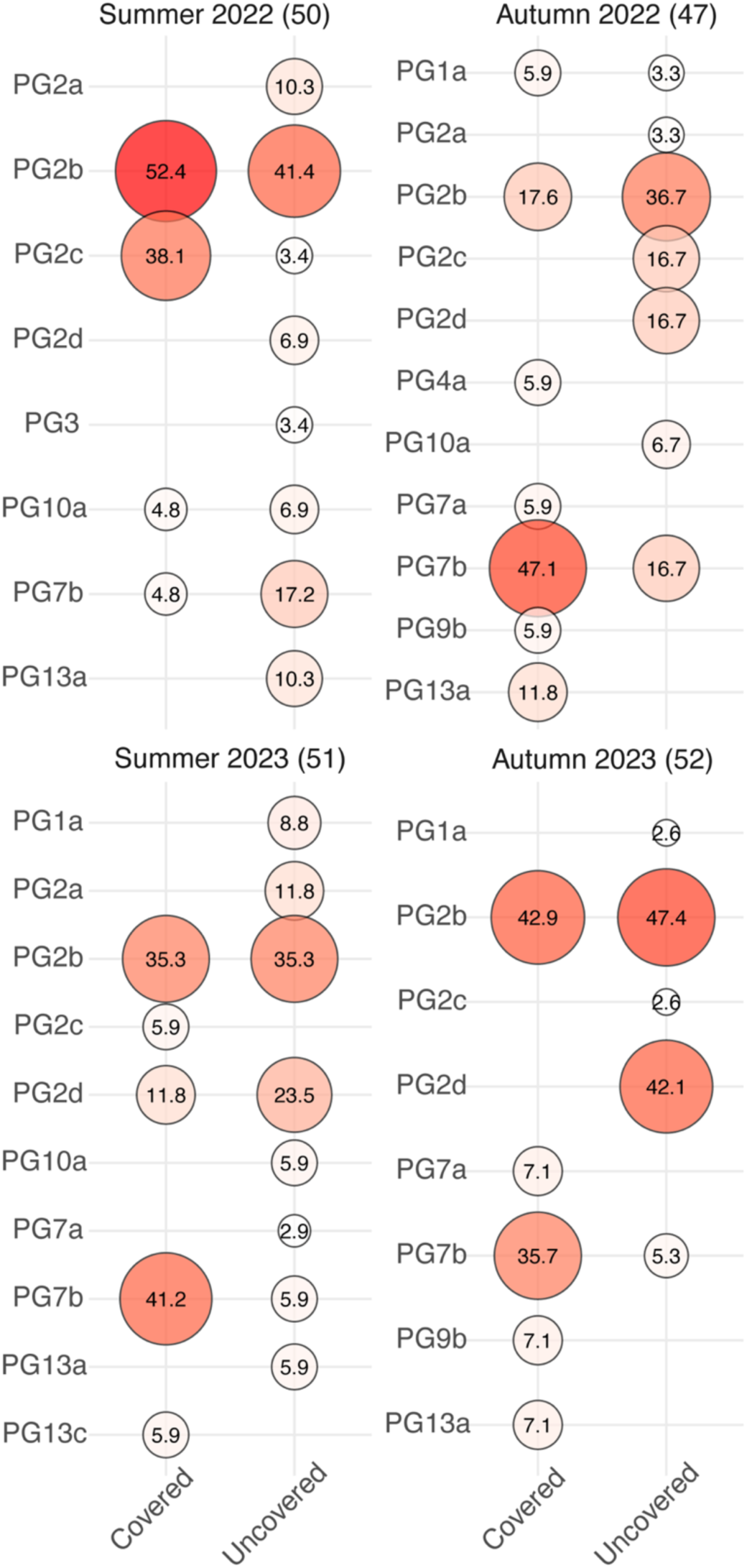
Percentages of strains of each clade of *P. syringae* (96% of ANI) isolated from covered or uncovered trees in individual samplings. The numbers in the circles show the percentages that each clade takes up from the total numbers of strains isolated from covered or uncovered trees in each sampling. The size and intensity of the colour of circles visualise the percentages in each circle. The total numbers of strains isolated from each sampling are marked in brackets next to the sampling time. Forty-two *Pss*9644-like strains (ANI ≥ 99.95%), progeny of inoculum, were excluded.

**Figure S3.**
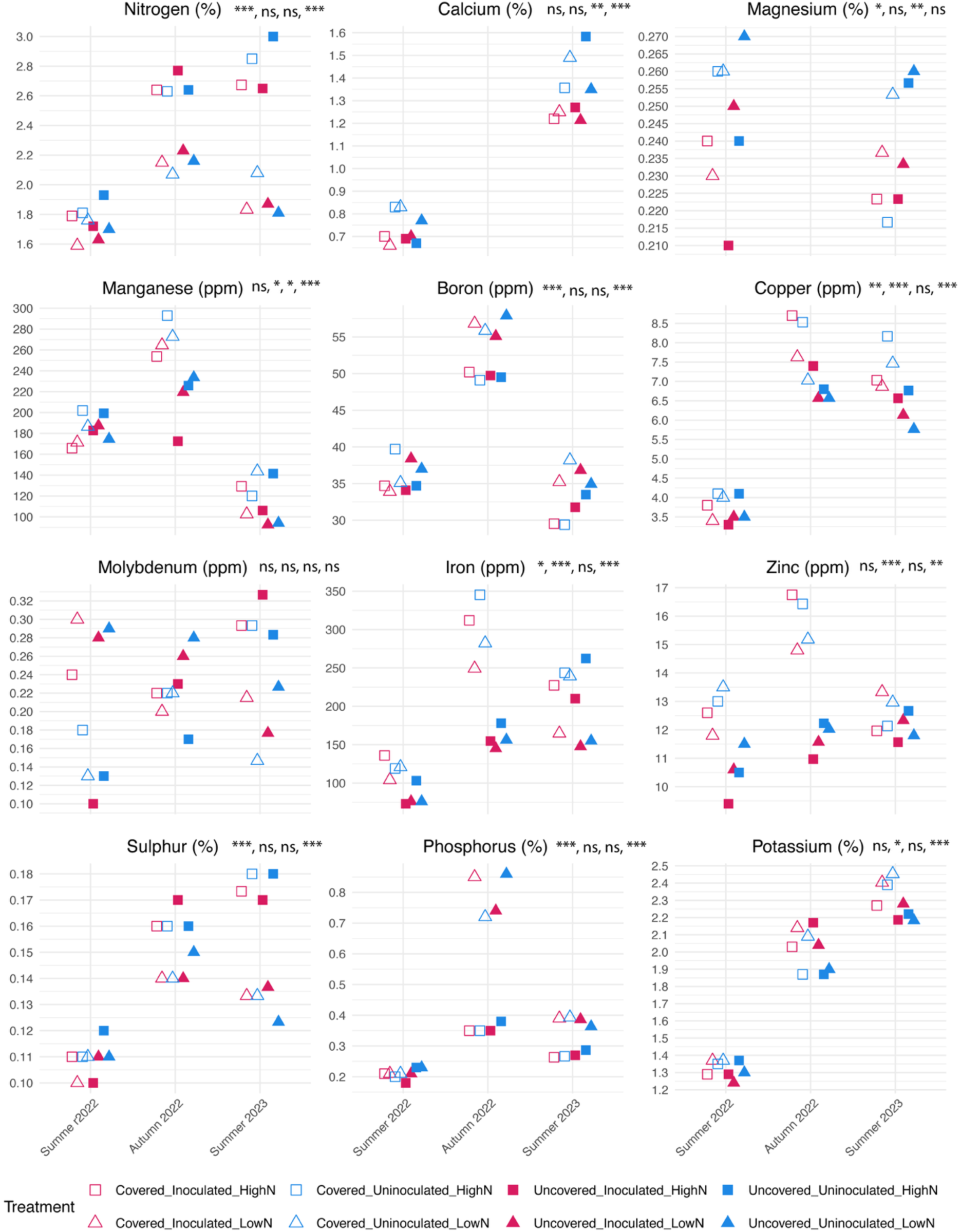
Nutrient levels of cherry leaves from different groups at different timepoints. Leaf samples from three blocks were collected and pooled together. Measurement and analysis were performed in Lancrop Laboratories. Measurement of summer 2023 is not available. Data of calcium and magnesium were missing due to technical issue. Differences in leaf nutrient concentrations were analysed using linear models (ANOVA) with fertigation, cover, inoculation and time as fixed factors. Significance levels for nitrogen fertigation, cover, inoculation and sampling time are indicated next to the title of each sub-panel, with ***p < 0.001, **p < 0.01, *p < 0.05, and ns denoting not significant.

**Figure S4.**
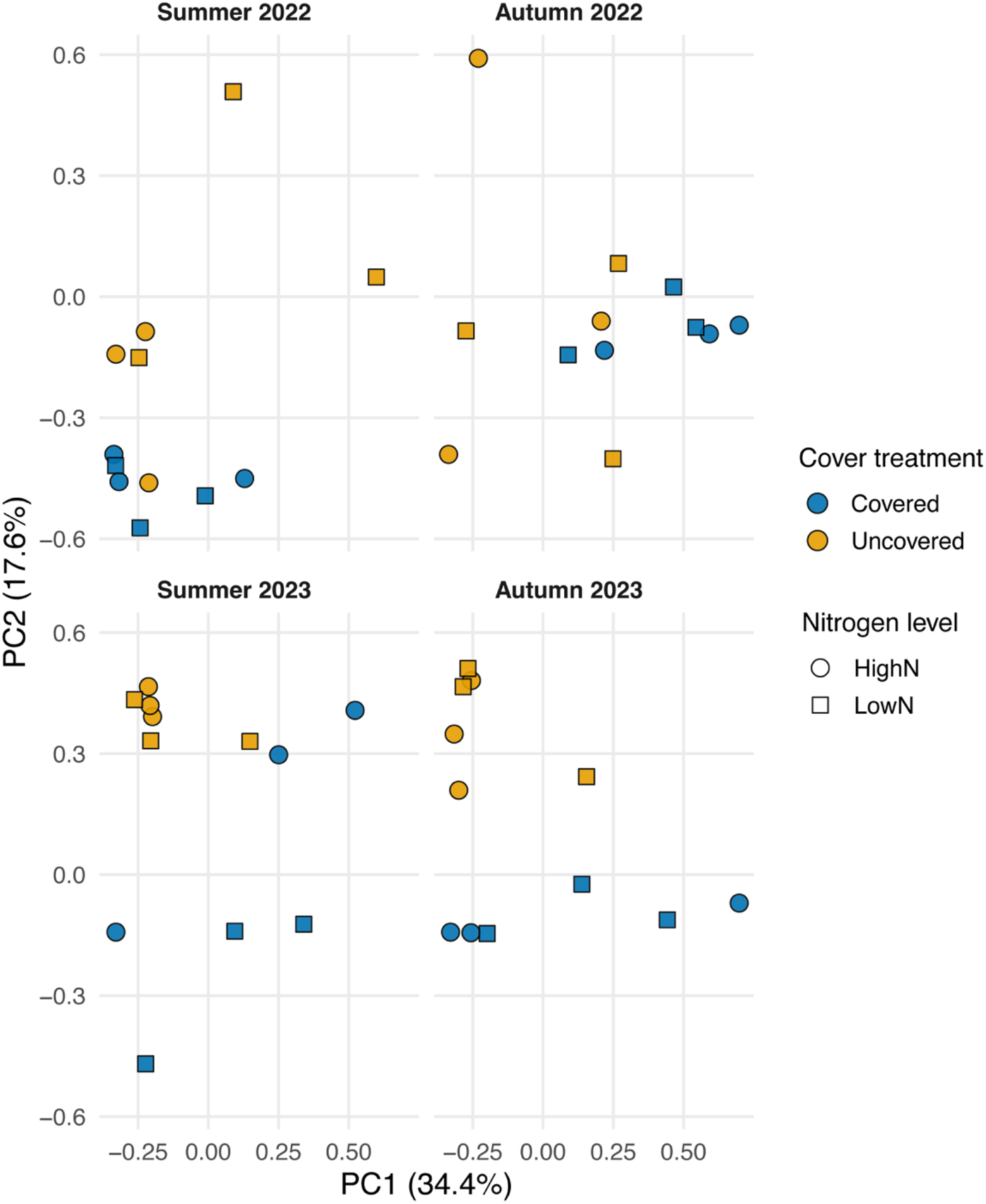
Principal component analysis of *P. syringae* community composition under combined nitrogen and cover treatments. The analysis was based on Hellinger-transformed community data and illustrates differences in *Ps* composition among cover and nitrogen treatments across four sampling timepoints. Points represent individual replicate blocks of each treatment. Symbols show nitrogen level (circles, high level of nitrogen fertigation; square, low level of nitrogen fertigation), and fill colours represent cover treatment (blue, covered; orange, uncovered). Each panel corresponds to a sampling timepoint. PC1 and PC2 explain 34.5% and 17.5% of the total variation, respectively. Forty-two *Ps*s9644-like strains (ANI ≥ 99.95%), progeny of inoculum, were excluded.

**Figure S5.**
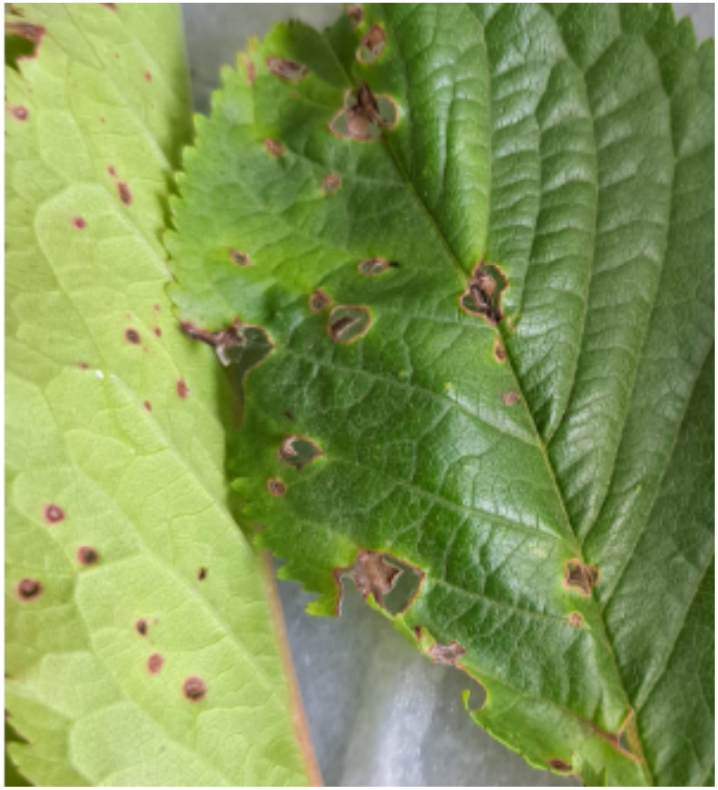
Shothole symptoms on cherry leaves associated with bacterial canker. Cherry leaves showing characteristic shothole symptoms. Necrotic lesions on the leaf surface eventually dry and fall out, leaving circular holes typical of bacterial shot-hole disease symptoms observed in infected trees.

**Figure S6.**
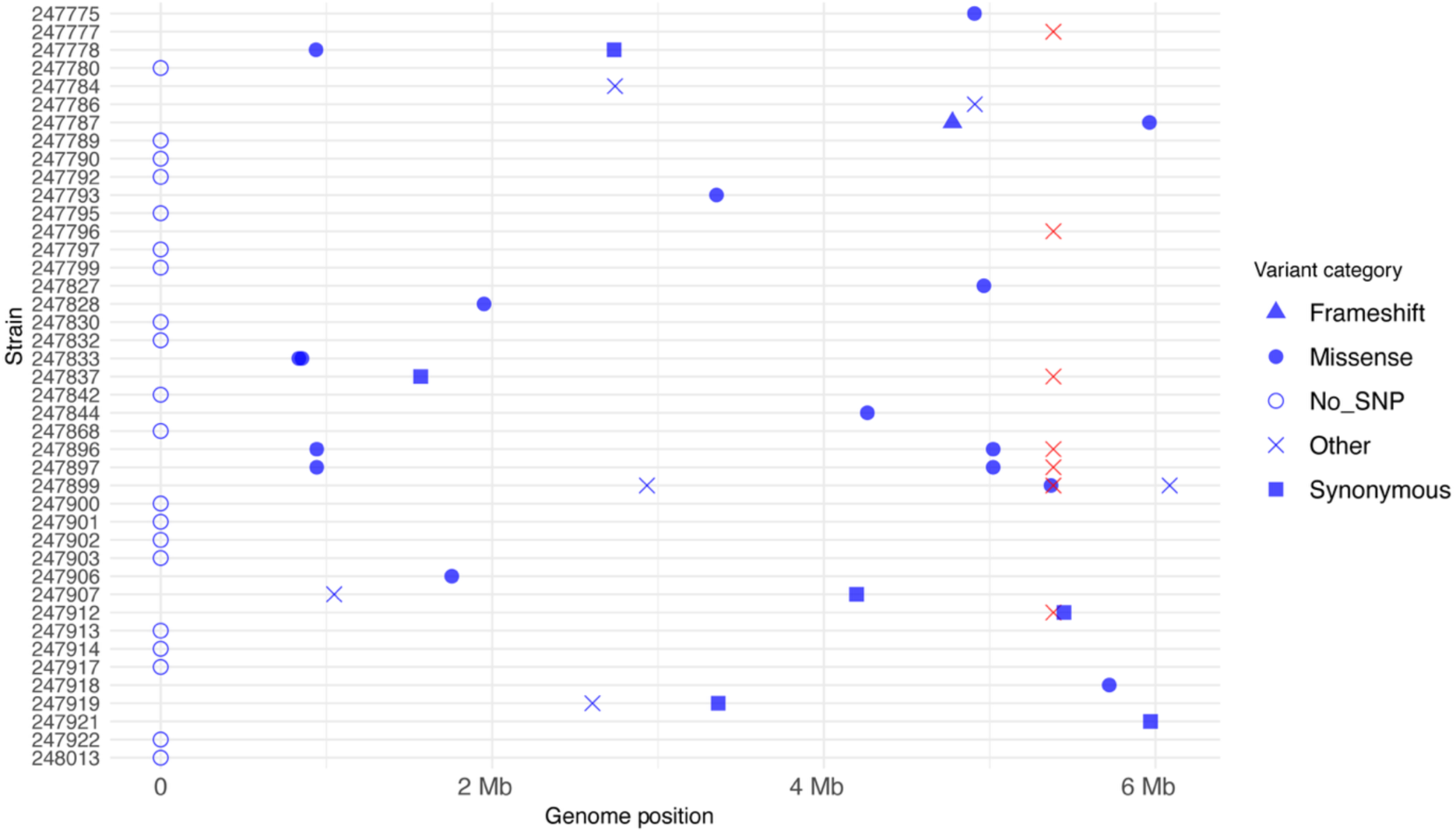
Genome-wide distribution of single-nucleotide polymorphisms (SNPs) identified in 22 re-isolated strains of *P. syringae* pv. *syringae* 9644 (*Pss*9644). SNPs were detected using Snippy, with the complete long-read reference genome of *Pss*9644 (GenBank accession GCA_023277945.1) as the reference. Each point represents a variant position detected in an individual strain, with shapes denoting variant type (missense, synonymous, frameshift, gene fusion or other). The direct-repeat deletions are highlighted in red.

**Figure S7.**
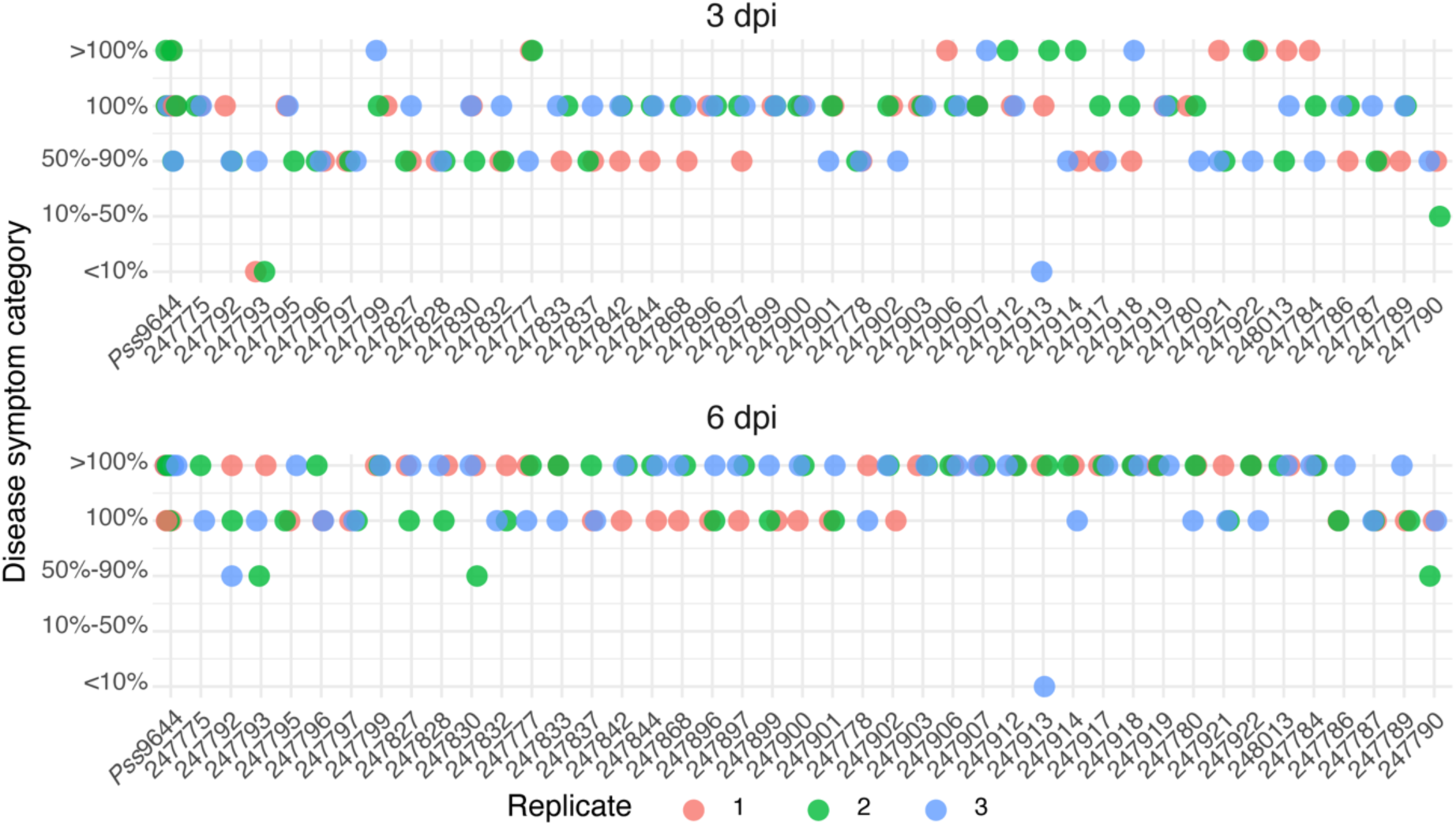
Virulence of re-isolated *P. syringae* pv. *syringae* 9644 (*Pss*9644) strains on cherry leaves. Leaf infiltration assays were conducted on detached leaves of *Prunus avium* cv. Sweetheart to assess lesion formation and discolouration symptoms. Symptom severity was categorised assessed based on the percentage browning /blackening at the inoculation site: No symptom, <10%, 10%–50%, 50%–90%, 100% discolouration and symptoms spreading from the infiltrated area (>100%). Bacterial overnight cultures (2×10^8^ CFU ml^-1^) were used as inoculum. Disease symptoms were scored 3– and 6-days post-inoculation.

**Figure S8.**
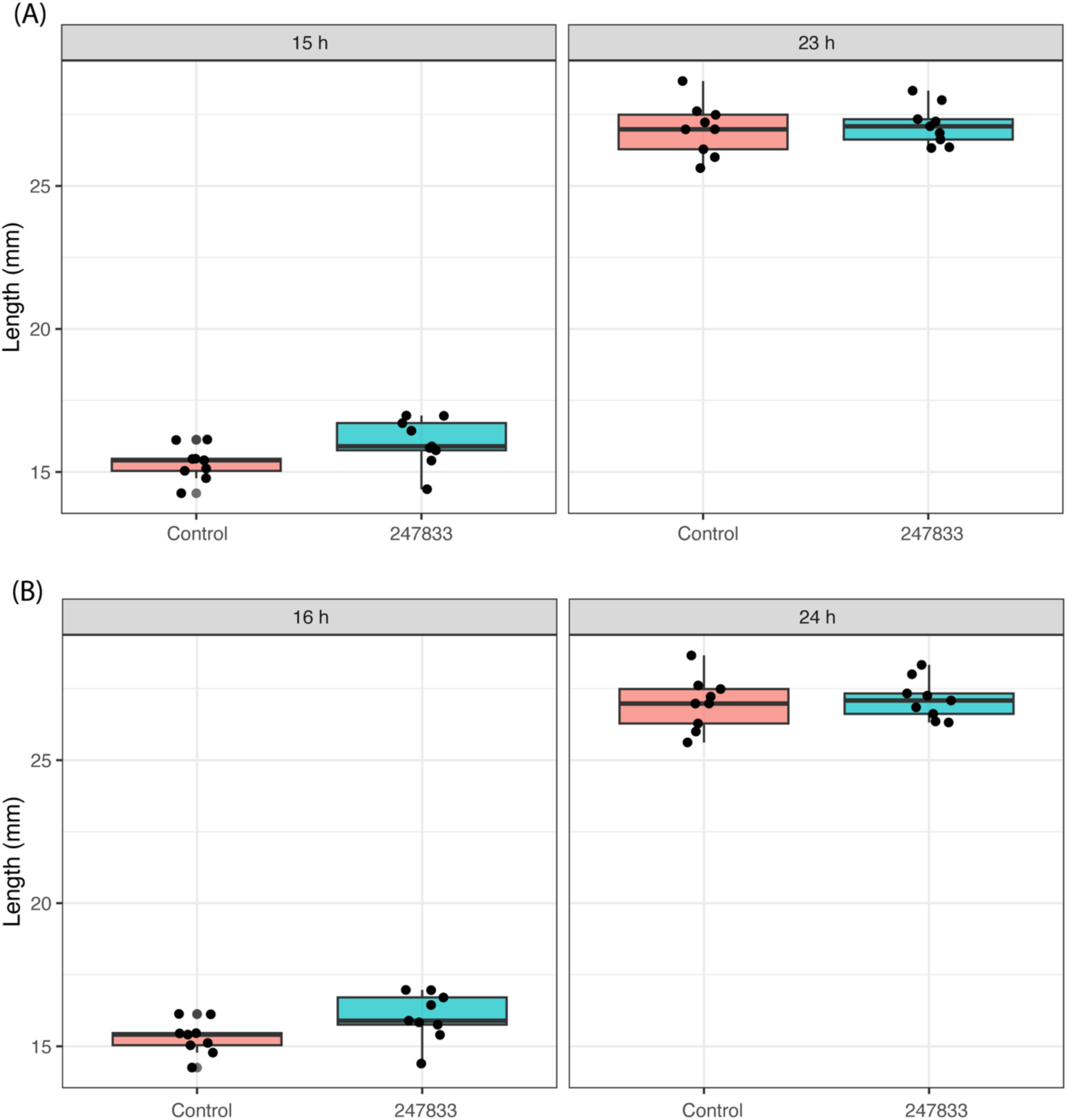
Swimming and swarming motility of *P. syringae* strains. **(A)** Swimming motility. **(B)** Swarming motility. Motility is shown as radial expansion from the inoculation point. The control strain (247792), a recovered *Pss*9644 isolate with no SNP detected, exhibited typical motility. No swimming or swarming was observed for the non-motile negative control strain *Pseudomonas savastanoi pv. fraxini* 1006.

**Table S1.**
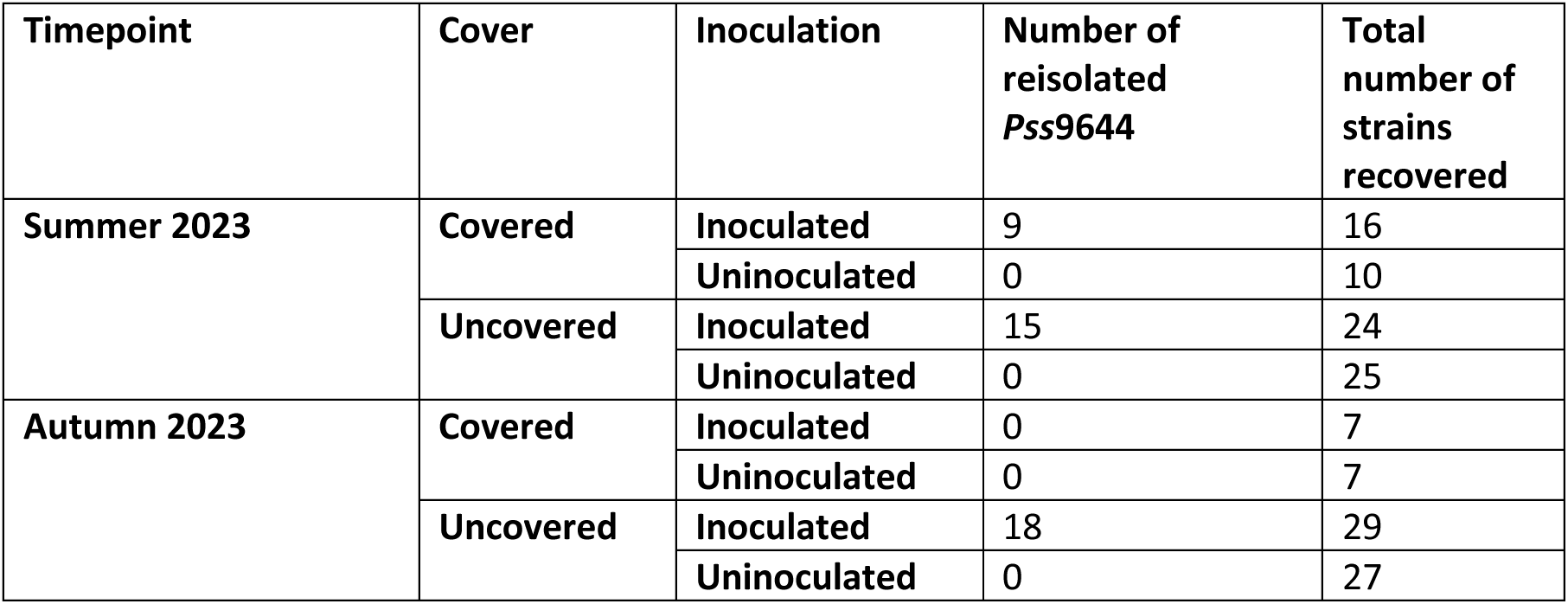
Number of strains recovered from different treatments and the numbers of reisolated *Pss*9644. Inoculation of trees occurred in January 2023. *Pss*9644 strains were differentiated by sharing over 99.95% ANI with the inoculated *Pss*9644.

| Timepoint | Cover | Inoculation | Number of reisolated *Pss9644* | Total number of strains recovered |
| --- | --- | --- | --- | --- |
| Summer 2023 | Covered | Inoculated | 9 | 16 |
|  |  | Uninoculated | 0 | 10 |
|  | Uncovered | Inoculated | 15 | 24 |
|  |  | Uninoculated | 0 | 25 |
| Autumn 2023 | Covered | Inoculated | 0 | 7 |
|  |  | Uninoculated | 0 | 7 |
|  | Uncovered | Inoculated | 18 | 29 |
|  |  | Uninoculated | 0 | 27 |

**Table S2.**
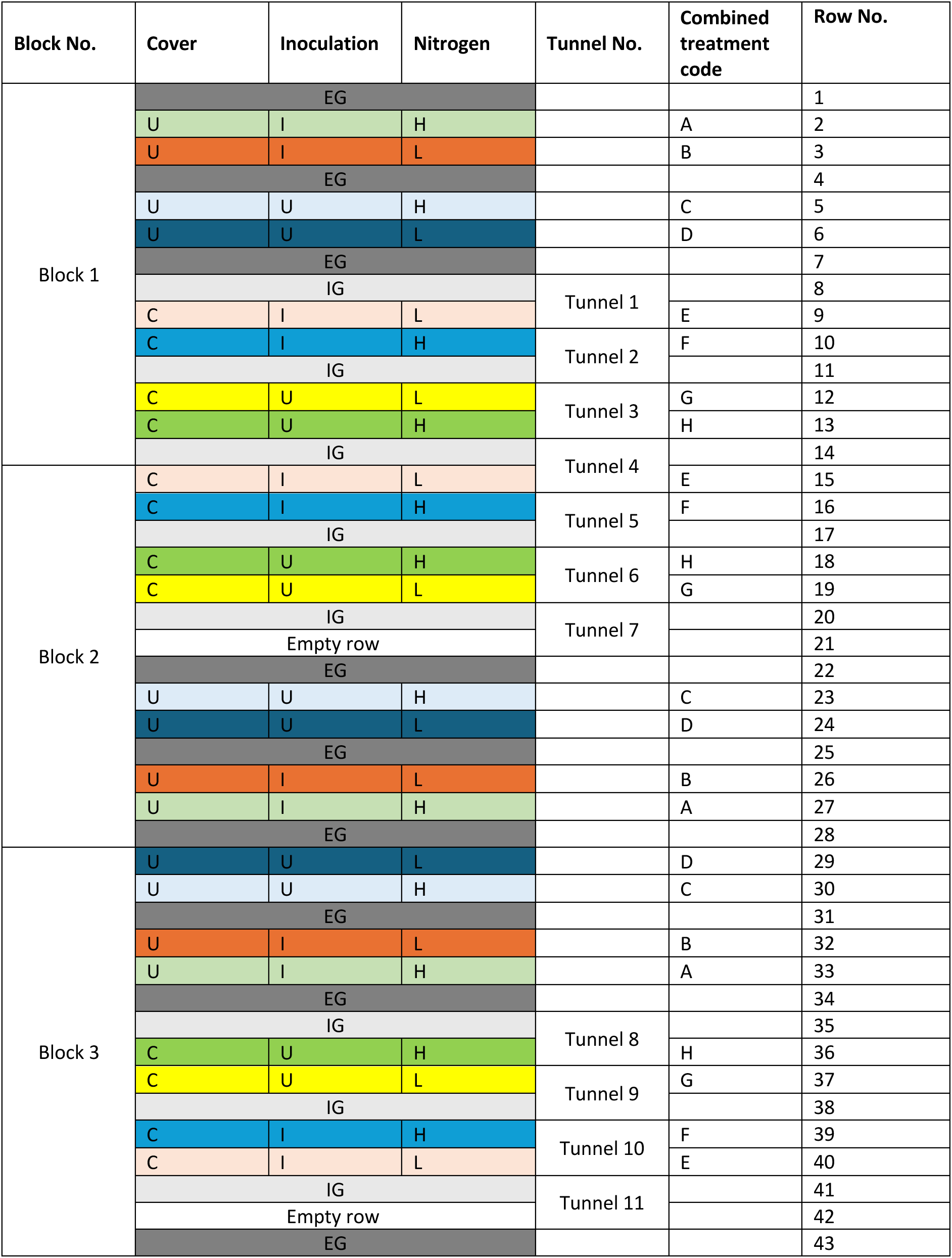
Layout of cherry trees under different treatments and guard trees in the split-plot field experiment at NIAB East Malling, UK. The main plot factor was polytunnel cover (C, covered; U, uncovered), and subplots were defined by nitrogen level (H, high; L, low) and pathogen inoculation (I, inoculated; U, uninoculated). Each combined treatment (A–H) represents one of eight treatment combinations and is colour-coded accordingly. Two rows of trees were placed under each polytunnel, and treatments were randomised and replicated across three blocks. EG, external guard trees located outside tunnels; IG, internal guard trees within tunnels separating different treatment subplots.

**Table S3.** Composition of nutrient solutions used for fertigation under low and high nitrogen treatments in 2022 and 2023. The fertigation formulation was revised in 2023 to enhance the difference in nitrogen concentrations between treatments.

| Nutrient (ppm) | 2022 |  | 2023 |  |
| --- | --- | --- | --- | --- |
|  | Low N | High N | Low N | High N |
| N-NO<sub>3</sub> | 0 | 34 | 21.4 | 154.1 |
| N-NH<sub>4</sub> | 60 | 96 | 2.6 | 11 |
| P | 212.5 | 212.9 | 73 | 73 |
| K | 98.8 | 97.4 | 300 | 300.5 |
| Ca | 15 | 15 | 60 | 60.1 |
| Mg | 0 | 0 | 44 | 44 |
| Na | 13 | 13 | 15.8 | 15.8 |
| S-SO<sub>4</sub> | 23 | 23 | 170.8 | 46.7 |
| Cl | 45 | 45 | 78.2 | 17 |
| Fe | 0 | 0 | 2.97 | 2.97 |
| B | 0 | 0 | 0.1 | 0.1 |
| Cu | 0 | 0 | 0.1 | 0.1 |
| Zn | 0 | 0 | 0.61 | 0.61 |
| Mn | 0 | 0 | 1.44 | 1.44 |
| Mo | 0 | 0 | 0.04 | 0.04 |
| EC (mS/cm) | 0.96 | 1.2 | 1.63 | 1.69 |

**Table S4.** Numbers of strains isolated/sequenced/confirmed to be *P. syringae* based on genomic sequences. Four strains were isolated from the two leaves collected from each tree sampled. To avoid sequencing clonal strains from the same tree, only one PCR-confirmed *Ps* strain from each tree was selected for whole genome sequencing.

| Treatment | Sampling timepoint |  |  |  | Total |
| --- | --- | --- | --- | --- | --- |
|  | Summer 2022 | Autumn 2022 | Summer 2023 | Autumn 2023 |  |
| Covered_Inoculated_HighN | 60/15/5 | 60/4/3 | 60/14/9 | 60/10/0 | **240/43/17** |
| Covered_Inoculated_LowN | 60/11/8 | 60/12/7 | 60/9/7 | 60/10/7 | **240/42/29** |
| Covered_Uninoculated_HighN | 60/11/6 | 60/9/3 | 60/14/3 | 60/15/6 | **240/49/18** |
| Covered_Uninoculated_LowN | 60/9/2 | 60/11/4 | 60/11/7 | 60/7/1 | **240/38/14** |
| Uncovered_Inoculated_HighN | 60/6/2 | 60/10/9 | 60/16/13 | 60/15/15 | **240/47/39** |
| Uncovered_Inoculated_LowN | 60/14/12 | 60/12/9 | 60/13/11 | 60/15/14 | **240/54/46** |
| Uncovered_Uninoculated_HighN | 60/8/8 | 60/10/7 | 60/13/13 | 60/15/14 | **240/46/42** |
| Uncovered_Uninoculated_LowN | 60/11/7 | 60/12/5 | 60/13/12 | 60/14/13 | **240/50/37** |
| **Total** | **480/85/50** | **480/80/47** | **480/103/75** | **480/101/70** | **1920/369/242** |

**Table S5.** Bacterial strains used in this study. [Note: An Excel file is submitted separately].

